# Structural and dynamic studies uncover a distinct allosteric modulatory site at the μ-opioid receptor

**DOI:** 10.1101/2025.10.14.682454

**Authors:** Haonan Zhang, Kirill Konovalov, Alexandra K. Parpounas, Davide Provasi, Shifan Yang, Alejandro Abraham, Audrey L. Warren, Gregory Zilberg, Suri Wang, Marta Filizola, Daniel Wacker

## Abstract

Positive allosteric modulators (PAMs) of the μ opioid receptor (MOR) offer a promising path toward safer opioid therapeutics, yet their mechanisms of action remain poorly understood. Here, we uncover the structural and mechanistic basis of BMS-986187, a chemically distinct MOR PAM with *in vivo* efficacy, using an integrated approach combining cryogenic electron microscopy (cryo-EM), molecular dynamics (MD) simulations, signaling assays, and site-directed mutagenesis. We identify a previously uncharacterized allosteric site for BMS-986187, a lipid-facing pocket formed by MOR transmembrane helices 2, 3, and 4, distinct from sites occupied by other known MOR PAMs or negative allosteric modulators. BMS-986187 engages both receptor residues and a neighboring cholesterol molecule, suggesting a cooperative ligand–lipid mechanism. Our studies pinpoint residues essential for allosteric modulation, while information-theory analysis of MD trajectories uncovers specific allosteric communication pathways linking the PAM site to both the orthosteric agonist DAMGO and the G protein interface. Together, these findings redefine the landscape of MOR allosteric modulation by revealing a novel binding site, a potentially lipid-sensitive allosteric mechanism, and the molecular wiring of long-range communication within MOR. This work provides a new molecular framework for the rational design of PAMs targeting opioid receptors with improved precision and possible therapeutic potential.

## Introduction

Opioids remain among the most effective painkillers available today, notwithstanding the numerous debilitating adverse effects associated with their prolonged use. Those include physical dependence, withdrawal, addiction, and opioid use disorder (OUD), a chronic, relapsing condition linked to high morbidity and mortality rates^1^. Opioid overdoses, often resulting from the misuse of prescription opioids (e.g., oxycodone, morphine) or unintentional exposure to highly potent synthetic opioids (e.g., fentanyl) frequently found adulterating recreational drugs (e.g., heroin), continue to pose a significant public health concern in the United States. The substantial limitations of current FDA-approved medications for OUD and opioid overdose, along with the chronic nature and high relapse risks associated with OUD, underscore the urgent need for developing more effective, patient-specific medications.

The µ opioid receptor (MOR) is the principal molecular target mediating both the therapeutic and adverse effects of current opioid medications^2^. Efforts to develop novel chemical entities targeting this receptor has generally followed three main strategies based on receptor interaction sites: (i) orthosteric modulators that bind to the classical, highly conserved opioid binding site utilized by endogenous ligands; (ii) allosteric modulators that target alternative binding sites distinct from the primary conserved region; and (iii) bitopic ligands that simultaneously interact with both orthosteric and allosteric sites^3^.

Over the past 20 years, most efforts have focused on developing novel orthosteric MOR agonists engineered for biasing MOR signaling, preferentially activating signaling pathways associated with analgesia while minimizing those responsible for adverse effects^4^. While this so-called “biased agonism” approach has garnered significant attention, it has thus far yielded underwhelming results in clinical trials^5^. Moreover, recent studies have questioned whether the only opioid as advanced under this strategy can truly be considered “biased”^6^. In comparison to biased agonism, allosteric modulation of MOR (and other GPCRs) is even less understood. To date, no MOR allosteric modulators have entered clinical use, and research into these compounds remains at an early stage, both at the mechanistic and pre-clinical levels. Nonetheless, recent studies have identified several chemical scaffolds with encouraging pharmacological activity *in vitro* and *in vivo*. For instance, DNA-encoded library screening has yielded a novel MOR negative allosteric modulator (NAM), compound 368^7^, while studies by our group^8^ and others^9,10^ have led to the discovery of chemically distinct MOR positive allosteric modulators (PAMs) with demonstrated *in vivo* efficacy and probe dependence, such as BMS-986122, BMS- 986187, MS1, and Comp5^8–10^. Despite these advances, our molecular understanding of how allosteric modulators influence MOR-mediated signaling remains limited.

Recent structural studies have revealed distinct binding sites for the MOR NAM compound 368^7^ and the PAM BMS-986122^11^. While compound 368 binds within the helical bundle of MOR, adjacent to the orthosteric binding site occupied by the small-molecule antagonist naloxone, BMS-986122 occupies a lipid-facing site located between transmembrane (TM) helices 3, 4, and 5 in MOR bound to the peptide agonist DAMGO^11^. However, in the absence of structural data for other PAMs, it remains unclear whether they engage the same binding site, as some have proposed^12,13^, or target distinct, uncharacterized allosteric sites. Moreover, little is known about how PAMs influence MOR conformational dynamics to elicit potential pathway-specific effects. It also remains to be determined whether structurally diverse PAMs act via shared or divergent mechanisms independent of the orthosteric ligand, despite functional evidence suggesting that MOR PAMs BMS-986122 and BMS-986187 may engage overlapping binding sites and modulate MOR signaling through conserved pathways^12,13^. Understanding these mechanisms is particularly important for evaluating potential synergistic interactions between PAMs that could amplify MOR signaling.

To begin addressing these questions, we herein combine cryogenic electron microscopy (cryo-EM), molecular dynamics (MD) simulations, signaling assays, and site-directed mutagenesis experiments. Our findings reveal that the PAM BMS-986187 binds to an allosteric site distinct from those previously reported. We propose a mechanism for its modulatory activity and provide structural and dynamic insights into how BMS-986187 may enhance MOR signaling.

## Results

### Identification of a unique BMS-986187 binding site at MOR

To investigate the mode of binding of BMS-986187 at MOR, we determined a cryo-EM structure of the MOR-Gα_i1_β_1_γ_2_ signaling complex in the presence of DAMGO and 100 µM BMS-986187 (Fig. 1A, Extended Data Figs. 1, 2, Extended Data Table 1). This was achieved by adapting previously established methods^14–16^ and co-expressing human MOR carrying a stabilizing F158^3.41^W mutation^14,17^ (superscripts denote generic GPCR numbering^18,19^) with an engineered heterotrimeric G protein. Specifically, Gγ_2_ was fused to Gα_i1_ and mutations were introduced into Gα_i1_ to strengthen interactions with Gβ_1_^15,16^.

**Fig 1.**
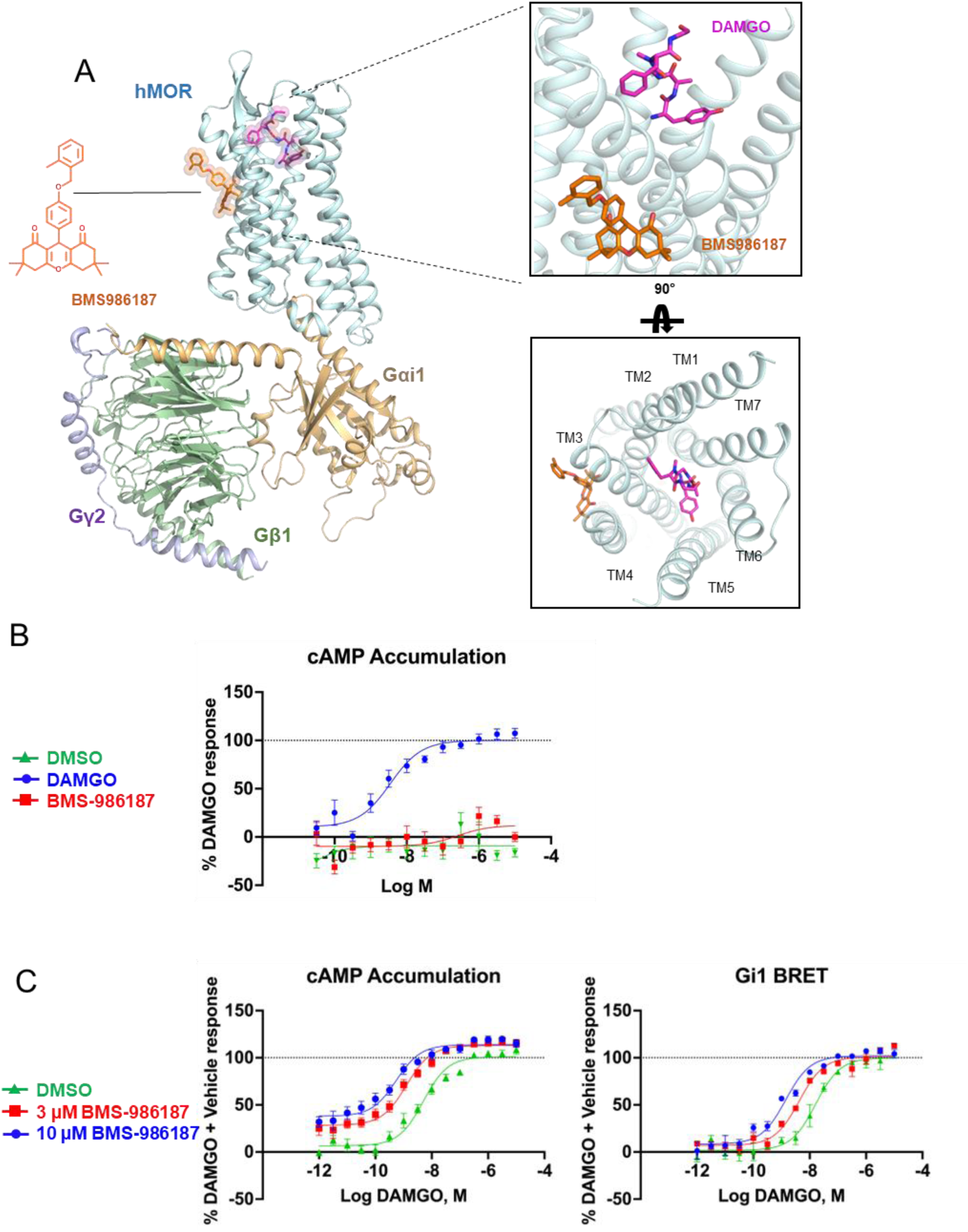
Pharmacological and structural characterization of BMS-986187 signaling at MOR. (A) overview of BMS-986187/DAMGO/MOR-Gi1 complex cryo-EM structure. Zoom-in shows BMS-986187 (orange) and DAMGO (magenta) bound at the receptor. MOR, Gαi1, Gβ1, and Gγ2, are shown in palecyan, wheat, green, and purple, respectively. (B) Intrinsic efficacy of BMS-986187, DAMGO, and vehicle (DMSO) at MOR as determined by cellular cAMP accumulation assay. (C) Allosteric effect of BMS-986187 on DAMGO signaling in both cAMP accumulation and Gi1 BRET assays. All data from signaling studies represent mean ± SEM of three to five independent experiments (n = 3-5) performed in triplicate, and have been normalized to the DAMGO + vehicle response.

Proteins were expressed in Sf9 insect cells, complex formation was induced by addition of DAMGO and BMS-986187, and samples were purified via affinity and size exclusion chromatography. We obtained a structure with an overall resolution of ∼3.5 Å, with local resolutions as high as 2.8 Å around the drug binding sites (Extended Data Fig. 1). We observe an antiparallel dimer arrangement with TM4 and TM5 helices at the interface, consistent with previous observations for MOR^14^ and other receptors^20^. While we obtained high-quality cryo-EM maps for MORs, the corresponding density for the G proteins was of lower resolution. Nevertheless, to provide a complete and biologically meaningful representation of the signaling complexes, we modeled the heterotrimers in their entirety rather than omitting select regions (see Methods for details).

The structure allowed us to unambiguously locate BMS-986187 in a membrane-facing crevice formed by TM2, TM3, and TM4 residues (Fig. 1A), which is distinct from the previously reported binding site of BMS-986122 (see subsection below). Given the unexpected binding location, we first sought to confirm previous pharmacological findings that classified BMS-986187 as an ago-PAM at MOR. To this end, we performed signaling assays in HEK293T cells transfected with human MOR and a cAMP biosensor we have used previously^15,16,21^. In brief, following stimulation with DAMGO and varying concentrations of BMS-986187 or vehicle, we added forskolin to elevate cellular cAMP levels and assessed MOR activation as a function of cAMP reduction. First, we assessed the intrinsic efficacy of BMS-986187 in this assay by performing concentration-response experiments in the absence of an orthosteric agonist (Fig. 1B). In our cAMP assay system, BMS-986187 did not exhibit notable intrinsic agonist activity at concentrations up to 10 µM. However, in the presence of the orthosteric agonist DAMGO, and consistent with earlier reports^9^, BMS-986187 produced a concentration-dependent amplification of DAMGO-induced signaling (Fig. 1C) characterized by a substantial increase in efficacy at low DAMGO concentrations and an increase in potency (Extended Data Table 2). The absence of intrinsic agonist activity of BMS-986187 at concentrations up to 10 µM indicates that the potentiation of DAMGO’s potency and efficacy is mediated by BMS- 986187 through allosteric mechanisms.

To validate these results, we conducted orthogonal assays using bioluminescence resonance energy transfer (BRET)-based G protein sensors to measure MOR signaling via G protein dissociation^22,23^. Using a G_i1_ sensor, we observed a concentration-dependent increase in DAMGO potency, with no significant effect on DAMGO efficacy (Fig. 1C). The differences observed between the BRET and cAMP assays likely reflect the distinct nature of these methods: BRET-based assays capture transient signaling events, whereas cAMP measurements are cumulative and therefore likely more sensitive to changes in maximal cAMP and signaling output.

### Characterization of BMS-986187’s binding mode at MOR

We next compared the BMS-986187 binding site to previously reported structures of MOR bound to allosteric modulators (Fig. 2). Although previous functional studies suggested that BMS-986122 and BMS-986187 might bind to the same site based on the absence of additive allosteric effects^12,13^, the structural data presented here provide the first direct evidence that these chemically distinct modulators occupy distinct binding sites. BMS- 986187 is positioned in a membrane-facing crevice formed by TM2, TM3, and TM4 on the outer leaflet of the membrane, whereas BMS-986122 has been shown by cryo-EM studies to bind a membrane-facing crevice formed by TM3, TM4, and TM5 on the inner leaflet of the membrane^11^. Both sites are different from the allosteric site reported for the NAM compound 368^7,11^, which binds near the orthosteric antagonist naloxone within an extended orthosteric binding pocket (Fig. 2).

**Fig 2.**
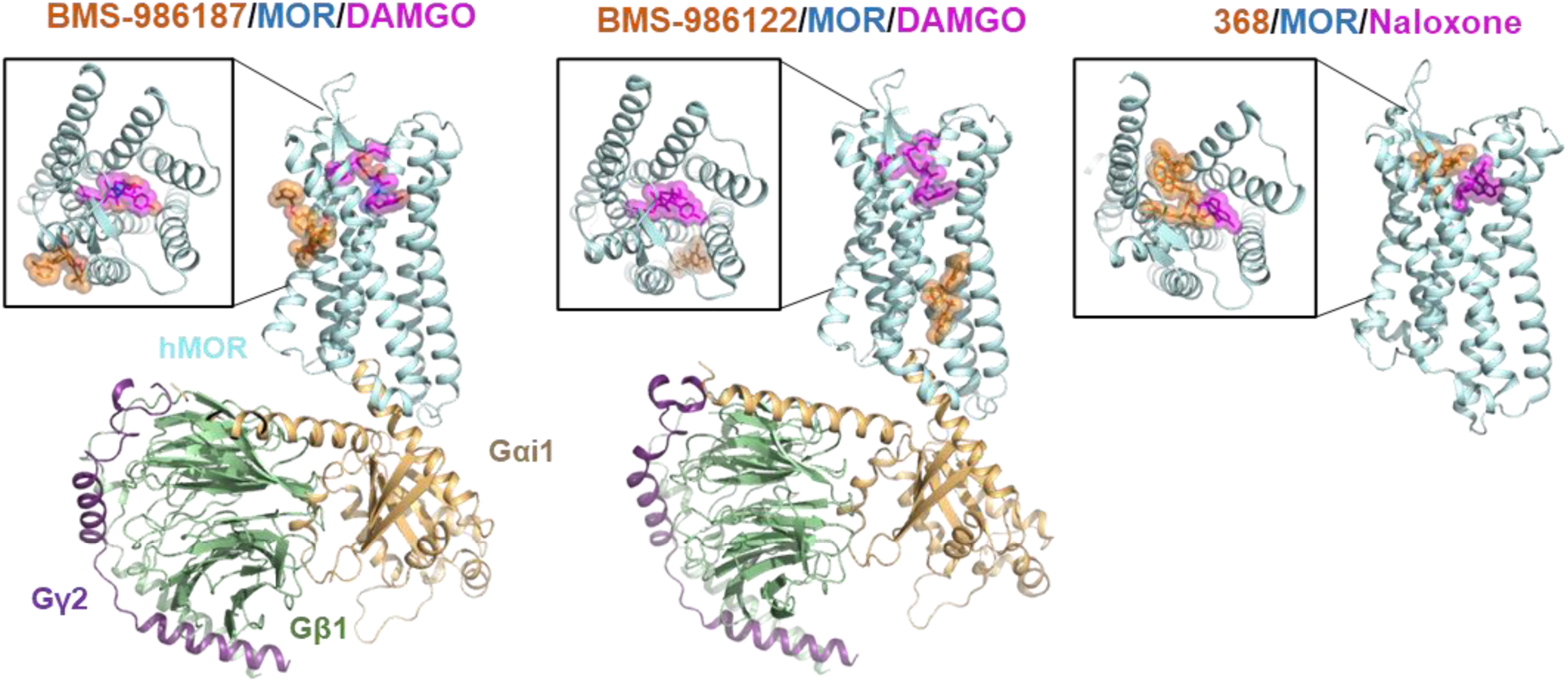
Binding location and pose of distinct allosteric modulators at MOR as determined by cryo-EM. Left, structure of the BMS-986187-bound DAMGO/MOR-Gi1 signaling complex (PDB: 9PU5). Middle, BMS-986122-bound DAMGO/MOR-Gi3 signaling complex (PDB: 8K9L). Right, compound 368-bound naloxone/MOR-Nb6 inactive-state complex (PDB: 9BJK). Zoom-ins represent extracellular views of the orthosteric binding pockets. PAMs, orthosteric ligands, MOR, Gαi1, Gβ1, and Gγ2, are shown in orange, magenta, palecyan, wheat, green, and purple, respectively.

Closer inspection of the BMS-986187 binding site reveals potential key interactions with receptor residues, as well as a nearby cholesterol (CLR) molecule (Fig. 3, Extended Data Fig. 2). To evaluate the stability of these BMS-986187 binding poses and their associated interactions, including those of DAMGO with the receptor, we carried out twenty independent 500-ns MD simulations of the BMS-986187-bound DAMGO/MOR-Gα_i1_β_1_γ_2_ signaling complex, totaling 10 μs, starting from a structure obtained through initial fitting to the cryo-EM density map prior to final refinement (Fig. 3, Extended Data Fig 3, Extended Data Table 3). As shown in Fig. 3A, most BMS-986187-receptor interactions inferred from the cryo-EM density (indicated by shaded bars) were retained throughout the simulations, with interaction frequencies exceeding 50% (compare full and shaded bars). Interactions were calculated between receptor residues and assigned fragments of BMS-986187 (see Methods), including: the ethyl-methyl-benzene (F1), the para-phenol (F2), the hexahydro-xanthene-dione (F3), the methylene group linking F1 and F2 (F4), and the bridging oxygen atom (F5). Among BMS-986187’s most stable interactions (>75% frequency), the van der Waals interaction with Y151^3.34^, observed consistently at 100% frequency via the F3 fragment, stands out. Additional frequent interactions included those with I144^3.27^ (via F1), I148^3.31^ (via F2), and M205^4.61^ (via F1, F2, and F3).

**Fig 3.**
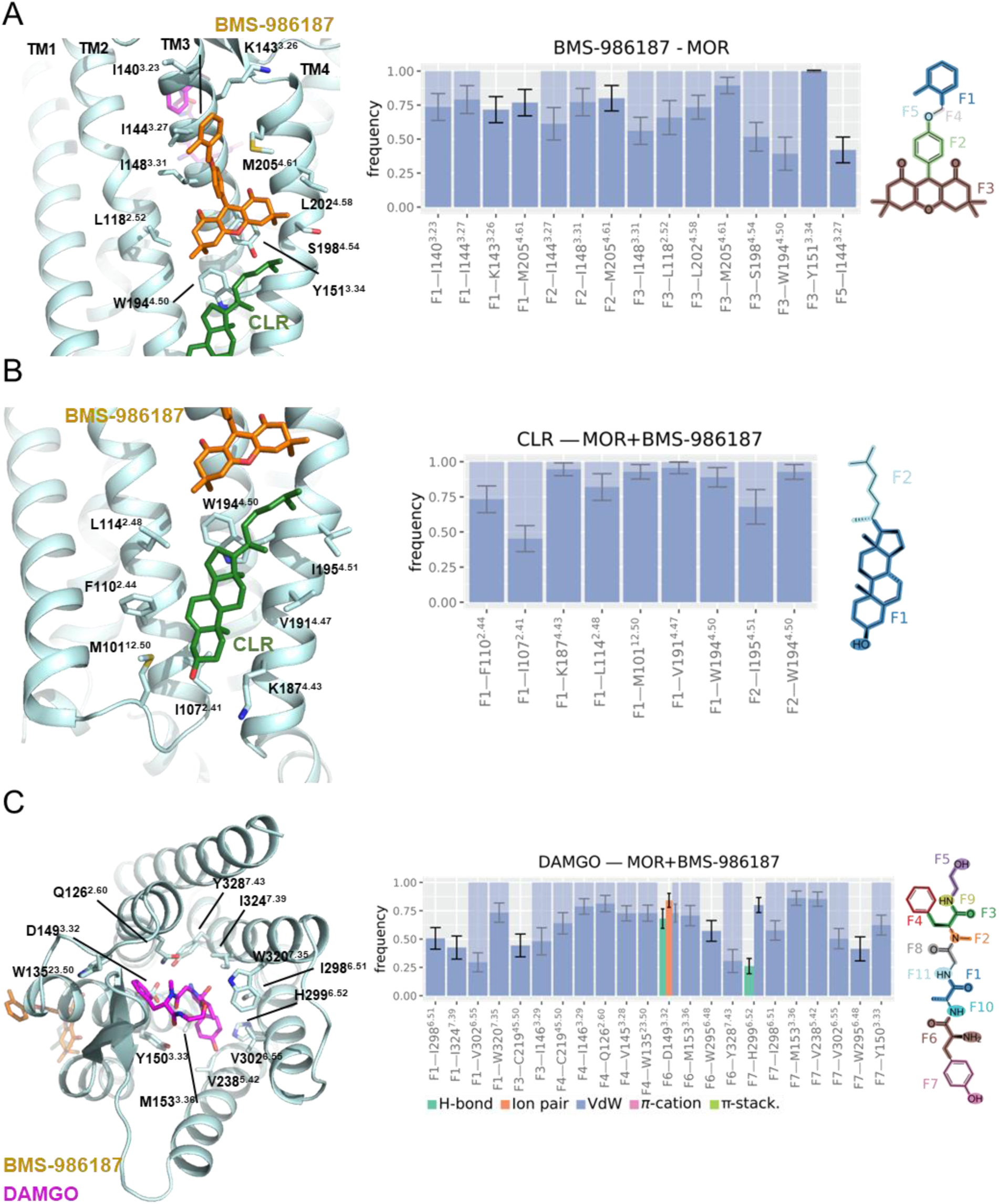
Ligand–MOR interactions observed in cryo-EM structures and MD simulations. (A) Interactions between BMS-986187 (orange), MOR (palecyan), and cholesterol (CLR, green). The left panel highlights interface residues identified in the cryo-EM structure. The middle panel shows the frequency of prevalent interactions (interaction frequency > 0.35) observed during MD simulations. Bar height reflects the mean interaction frequency across all simulation frames, with error bars representing the standard deviation computed over 25 ns time blocks. Blue shading over a bar indicates that the interaction is also observed in the cryo-EM structure. Interactions involving individual ligand atoms were grouped by molecular fragment, as illustrated in the right panel. (B) Interactions between MOR and a cholesterol molecule identified in the cryo-EM structure. (C) Interactions between DAMGO (magenta), bound in the orthosteric site, and MOR.

The largest decrease in interaction frequency was observed between the F3 fragment of BMS-986187 and W194^4.50^, which dropped to 39% during the simulation (Fig. 3A). Notably, W194^4.50^ simultaneously forms stable van der Waals interactions with both the sterol core (F1) and the aliphatic substituent (F2) of the nearby cholesterol molecule resolved in the cryo-EM map (Fig. 3B). This cholesterol molecule remains stably bound throughout the MD simulation, primarily through persistent (>75% of frames) van der Waals interactions with M101^12.50^ in intracellular loop 1 (IL1), as well as with L114^2.48^, K187^4.43^, and V191^4.47^, all mediated via its F1 fragment (Fig. 3B).

As expected for a flexible peptide, DAMGO exhibited greater conformational variability within the binding pocket compared to the more rigid BMS-986187 and cholesterol molecules, with several contacts observed in the cryo-EM structure maintained in fewer than 50% of simulation frames (compare full and shaded bars in Fig. 3C). Despite this flexibility, DAMGO maintained an overall structural stability throughout the simulations, with its most persistent interactions (>75% frequency) including van der Waals contacts between the phenyl F4 moiety and Q126^2.60^ and I146^3.29^, a salt-bridge between the F6 amine moiety and D149^3.32^, and van der Waals interactions between the F7 phenoxyl moiety and both M153^3.36^ and V238^5.42^. In contrast, the most affected interactions, though still occurring in >35% of frames, included contacts between the F1 moiety and V302^6.55^, and between the F6 moiety and Y328^7.43^ (Fig. 3C). However, these were compensated by new interactions formed between F1 and both I298^6.51^ and I324^7.39^ (compensating for the loss of interaction with V302^6.55^), and between F6 and W295^6.48^ (compensating for the loss of interaction with Y328^7.43^).

### Molecular mechanisms of BMS-986187

To elucidate the mechanistic basis by which BMS-986187 modulates MOR activity, we employed an integrated structural, dynamic, and functional approach. As an initial step toward understanding the conformational changes induced by BMS-986187, we determined a cryo-EM structure of the DAMGO/MOR-G_i1_ signaling complex in the absence of the PAM, using the same methodology as for the PAM-bound complex (Extended Data Fig. 3-4, Extended Data Table 1). Consistent with the PAM-bound complex, this structure also adopts an antiparallel dimer configuration, in agreement with previous observations by us and others^14,20^. The absence of BMS-986187 did not alter this arrangement, suggesting that the dimer interface is not influenced by the PAM. The BMS-986187-free structure was resolved at a global resolution of 2.89 Å, with local resolutions reaching ∼2.5 Å in the orthosteric and allosteric binding pockets, enabling high-confidence structural comparisons (Extended Data Fig. 4A, Extended Data Figure 3).

Structural comparison of the DAMGO/MOR-G_i1_ complexes with and without bound BMS- 986187 revealed minimal overall differences, with root-mean-square deviations (RMSDs) of 1.520 Å for the full complex and 0.538 Å for the receptor alone (Extended Data Fig. 4A). Despite this high degree of similarity, minor shifts in the positioning of the Gα_i1_ subunits were observed; however, limited G protein resolution precluded a detailed analysis of potential BMS-986187-mediated transducer rearrangements using cryo-EM data alone. At the BMS-986187 binding site, only subtle side chain reorientations were detected. However, a significant change was observed in the binding pose of the nearby cholesterol molecule, whose aliphatic tail and even sterol core shifts, likely to accommodate BMS-986187 in the PAM-bound complex (Extended Data Fig. 4A).

To assess the impact of BMS-986187 on the dynamics of the DAMGO/MOR–G_i1_ complex, particularly its effects on DAMGO-MOR and MOR-G protein interactions, we performed MD simulations of the complex without bound BMS-986187 (Extended Data Table 3), and compared the results to those of the PAM-bound system. DAMGO’s overall interaction pattern with MOR remained largely consistent regardless of PAM presence (Extended Data Fig. 4B); however, a subset of DAMGO-MOR interactions were notably affected by BMS-986187, including F3-C219^45.50^, F6-W295^6.48^, F7-I298^6.51^, and F1-V302^6.55^. Beyond DAMGO-MOR interactions, BMS-986187 also influenced the local membrane environment by altering the MOR–cholesterol interface. In the presence of the PAM, cholesterol lost its F2-I148^3.31^ interaction, exhibited a weakened F2-Y151^3.34^ interaction, and gained a markedly stronger F2-I195^4.51^ interaction (Extended Data Fig. 4B). These changes are consistent with the conformational shift in cholesterol’s binding pose observed between the PAM-bound and PAM–free DAMGO/MOR-G_i1_ complexes (Extended Data Fig. 4A). Finally, the MOR-G protein interface remained largely conserved regardless of BMS-986187 binding (Extended Data Fig. 4C). The most notable PAM-induced differences at the MOR–Gα_i1_ interface involved the disruption of key hydrogen bonds and salt bridges, specifically between MOR R265 and Gα_i1_ D341, and between MOR K273^6.26^ and Gα_i1_ E318.

To test whether specific MOR residues that directly interact with BMS-986187, based on cryo-EM density and confirmed by MD simulations (Fig. 3A), contribute to BMS-986187 binding and/or its allosteric modulation of DAMGO potency or efficacy, we performed cAMP signaling assays using mutant MOR constructs (Fig. 4, Extended Data Fig. 5, Extended Data Table 2). For each mutant, we collected DAMGO concentration-response data in the presence of vehicle (DMSO), 3 µM BMS-986187, and 10 µM BMS-986187. To facilitate interpretation, we plotted the data as bar graphs assessing the effects of mutations on: i) DAMGO potency (Fig. 4B), ii) BMS-986187-mediated potentiation of DAMGO potency (Fig. 4C), and iii) BMS-986187-mediated enhancement of DAMGO’s base efficacy (Fig. 4D) as outlined in Extended Data Fig. 5B.

**Fig 4.**
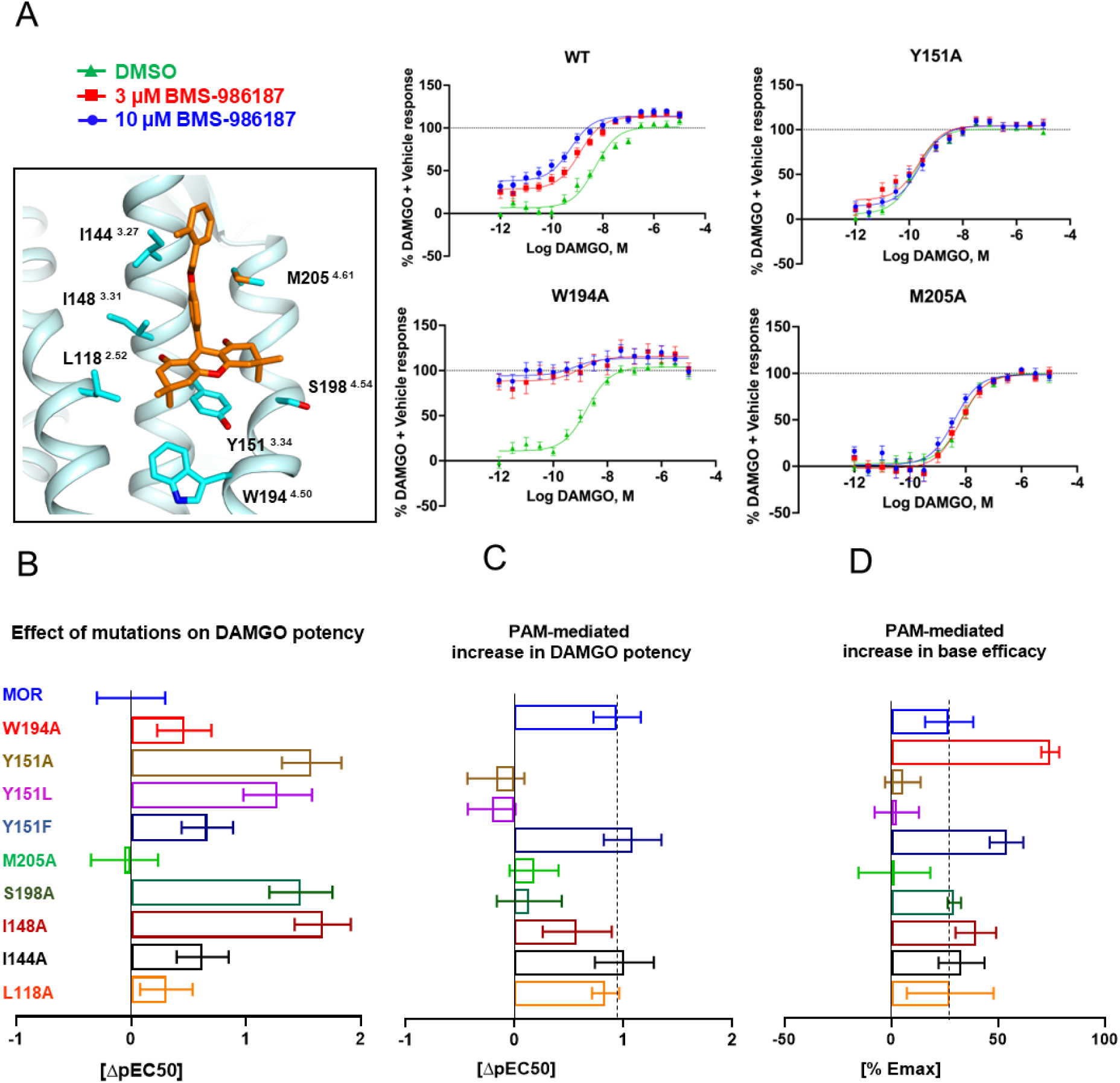
Site directed mutagenesis of BMS-986187-MOR interactions. (A) Binding site of BMS-986187 (orange), with MOR residues shown in cyan that were tested in cAMP accumulation assays. DAMGO concentration response data in the presence of vehicle (green), 3 μM BMS-986187 (red), and 10 μM BMS-986187 (blue) at wildtype and mutant MOR. Data represent mean ± SEM of three independent experiments (n = 3) performed in triplicate, and have been normalized to the DAMGO + vehicle response. (B) Bar graphs showing the effect of mutations on DAMGO potency, expressed as potency difference between mutant and wildtype MOR (ΔpEC50 = pEC50_Mut_ – pEC50_WT_). Larger numbers indicate that DAMGO exhibits increased potency at mutant construct. (C) Bar graph showing PAM-mediated increases in DAMGO potency, expressed as potency difference at each mutant or wildtype between vehicle (DMSO) and 10 μM BMS-986187 (ΔpEC50 = pEC50_BMS-986187_ – pEC50_DMSO_). Larger numbers indicate that BMS-986187 has an increased effect on DAMGO potency at the examined construct. (D) Bar graph showing PAM-mediated increases in DAMGO base efficacy (Emin), expressed as ΔEmin between vehicle (DMSO) and 10 μM BMS-986187 (ΔEmin = Emin_BMS-986187_ – Emin_DMSO_). Larger numbers indicate a more pronounced effect of BMS-986187 on Emin. Data represent mean ± SD calculated from three independent experiments (n = 3), and graphs for B, C, D were obtained by performing a multiple t test between mean values from individual experiments.

Of the ten MOR residues that maintain contact with BMS-986187 in more than 35% simulation frames (Fig. 3A), we mutated all except I140^3.23^, K143^3.26^, and L202^4.58^, which were excluded because mutations at these sites were expected to produce effects redundant with those of other targeted residues. As predicted by our structural and simulation data, several mutations altered the allosteric activity of BMS-986187, though the magnitude of these effects varied (Figs. 4B-D, Extended Data Fig. 5A). For example, mutations of Y151^3.34^, which forms van der Waals contacts with BMS-986187 in 100% of the simulation frames (Fig. 3A), were among the most impactful on the compound’s allosteric effects. The Y151^3.34^A mutation nearly abolished BMS-986187’s ability to enhance DAMGO’s efficacy at low concentrations (Fig 4A). We define this efficacy as DAMGO’s base efficacy (Emin) (Extended Data Fig 5B), as BMS-986187’s inability to increase this activity in the absence of DAMGO indicates that this does not reflect basal receptor activity. Compared to the effects on Emin, Y151^3.34^A appears to increase DAMGO potency in the absence of the PAM (Fig. 4B). Together, these data suggest that Y151^3.34^A may stabilize an active receptor conformation or enhance DAMGO-mediated activation independently of the PAM. To determine whether π-stacking or other van der

Waals interactions with Y151^3.34^ are critical for PAM activity, we also substituted this residue with phenylalanine and leucine. The Y151^3.34^F mutant preserved or slightly enhanced BMS-986187’s efficacy (Figs. 4C, 4D), whereas the Y151^3.34^L mutant completely abolished PAM activity (Figs. 4C, 4D), indicating that a π-stacking interaction between Y151^3.34^ and BMS-986187 might be important for allosteric modulation.

Another critical residue we tested is M205^4.61^, which forms stable interactions with multiple components of BMS-986187 (Fig. 3A) according to both our structure and simulations. Functional studies further support its importance: at the M205^4.61^A mutant, BMS-986187 loses its ability to modulate DAMGO activity (Figs. 4A, 4C, 4D), although DAMGO potency remains unchanged relative to the wild-type receptor (Fig. 4B). This suggests that while M205^4.61^ is not required for receptor activation by DAMGO, it is essential for mediating the allosteric effects of BMS-986187. We also examined residues L118^2.52^, I144^3.27^, I148^3.31^, and S198^4.54^, which were predicted to contribute to BMS-986187 binding and/or activity (Fig. 3A). Alanine substitutions at L118^2.52^, I144^3.27^, and I148^3.31^ had minimal impact on PAM activity. In contrast, the S198^4.54^A mutation had little effect on the increase in DAMGO’s Emin observed with 10 µM BMS-986187 (Fig. 4D), but abolished modulation at 3 µM (Extended Data Fig 5A). Moreover, the S198^4.54^A substitution eliminated BMS- 986187’s ability to enhance DAMGO potency (Fig. 4C). Interestingly, both I148^3.31^A and S198^4.54^A mutants led to increased DAMGO potency in the absence of PAM, suggesting that these residues may contribute to stabilizing the active conformation of MOR or specifically enhancing DAMGO-mediated receptor activation.

Although W194^4.50^ is centrally located within the PAM binding site, neither our structural data nor frequency interaction analysis from simulations support a primary role for this residue in mediating BMS-986187’s allosteric effects, as its contacts with the ligand are markedly reduced during simulations (Fig. 3A). Instead, W194^4.50^ consistently interacts with a nearby cholesterol molecule, regardless of BMS-986187’s presence (Fig. 3B, Extended Data Figs 4A-B). Unexpectedly, a W194^4.50^A mutation significantly enhanced the allosteric effects of BMS-986187, both by increasing its potentiation of DAMGO base efficacy (Fig. 4D) and by lowering the concentration required to achieve this effect (Fig. 4A).

Taken together, these findings underscore the contribution of specific MOR residues to both allosteric modulation and direct activation of the DAMGO/MOR-G_i1_ complex by BMS- 986187.

### State-specific conformations of key residues in the BMS-986187 binding pocket

We next examined the conformational states of key residues in the BMS-986187 binding pocket, specifically W194^4.50^ and Y151^3.34^, which have been functionally implicated in mediating BMS-986187’s allosteric activity (Fig. 4). To do so, we analyzed over 20 published high-resolution MOR structures^7,11,14,20,24–32^, including inactive states, a nanobody-stabilized active-like state, and several G protein-bound active states bound to agonists of varying efficacy (Fig. 5, Extended Data Fig. 6). Interestingly, we observe different rotameric conformations (as defined by the closest discrete rotamer in Coot^33^) of W194^4.50^ (m-95, t-90, t-105) and Y151^3.34^ (m-85, t-80, m -30) depending on MOR’s activation state, and seemingly independent of the bound orthosteric ligand (Fig. 5A, Extended Data Fig. 6). In our cryo-EM structures of the active DAMGO/MOR-G_i1_ complex, both with and without BMS-986187, as well as in all available agonist-bound MOR active structures, Y151^3.34^ adopts a t-80 rotamer pointing toward the cytosolic face of the receptor, while W194^4.50^ assumes an m-95 rotamer oriented parallel to the membrane (Fig. 5A, Extended Data Fig 6A). In contrast, in all available inactive MOR structures, Y151^3.34^ adopts an m-85 rotamer directed toward S198^4.54^ and the membrane, whereas W194^4.50^ (m-95) generally retains the same rotameric state observed in all MOR active structures. Notably, in the NAM-bound inactive structure (PDB: 9BJK), W194^4.50^ (t-105) and Y151^3.34^ (m -30) were modeled differently, though the cryo-EM density appears insufficient to unambiguously assign these states. As expected, the stable binding poses of BMS-986187 and the adjacent cholesterol molecule, both directly interacting with Y151^3.34^, constrain this residue to a trans rotamer oriented toward the cytosolic face of the receptor throughout the MD simulation. In the PAM-free DAMGO/MOR–G_i1_ simulations, where cholesterol is also present, Y151^3.34^ likewise favors the t-80 rotamer, although a small fraction (∼5%) of “inactive-like” m-85 states is observed. In contrast, W194^4.50^, despite reducing its interaction with BMS-986187 during simulation (Fig. 3A), consistently adopted an m-95 rotamer oriented parallel to the membrane in both PAM- bound and PAM-free DAMGO/MOR–G_i1_ simulations.

**Fig 5.**
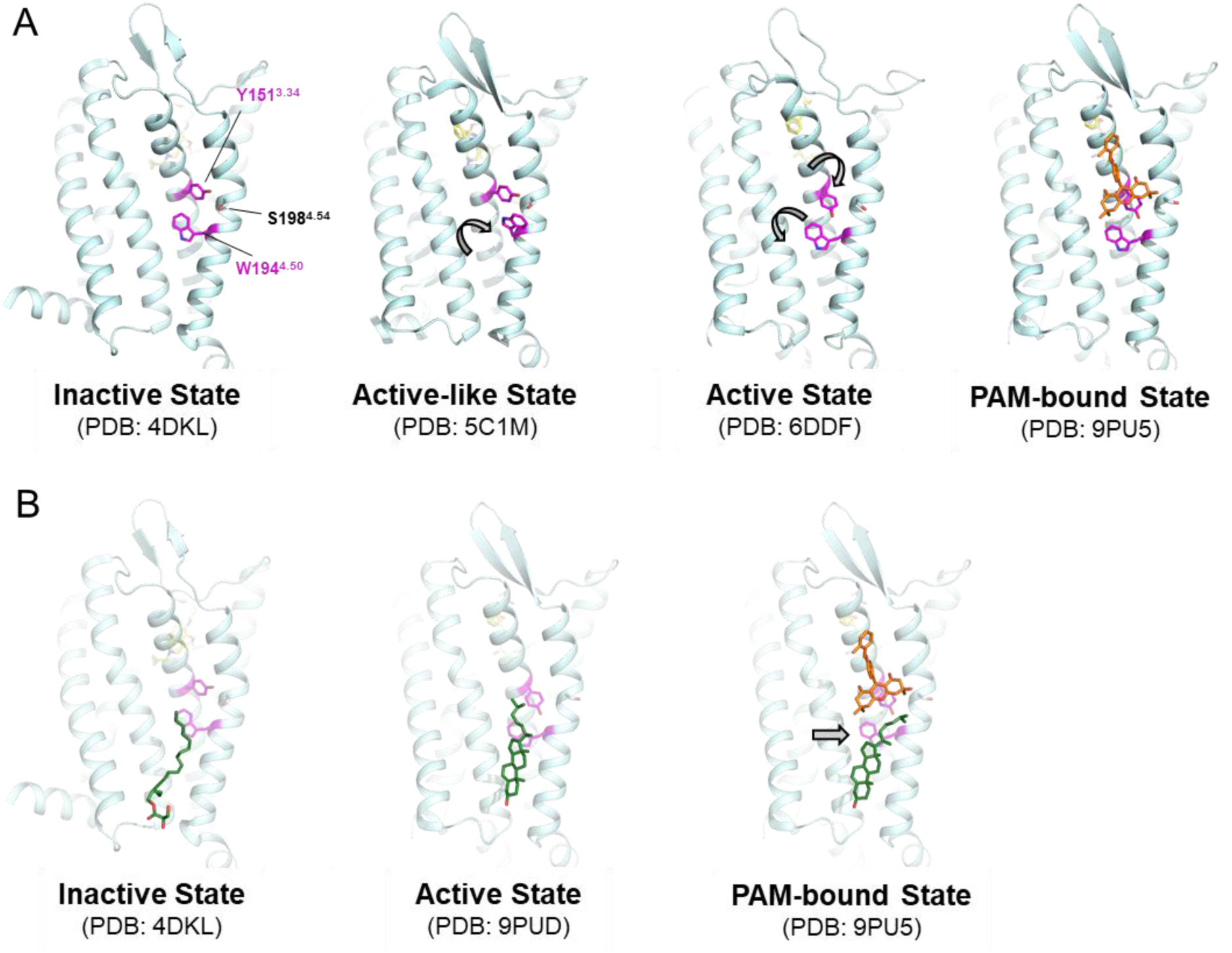
Structural analysis of Y151 and W194 conformations across different MOR activation states. (A) Representative structures of inactive- (PDB: 4DKL), active-like (PDB: 5C1M), active- (PDB: 6DDF) and BMS-986187-bound active-state (PDB: 9PU5) of MOR, with distinct rotamer configurations of Y151 and W194. (B) Representative structures highlight the binding of lipids and cholesterol near the BMS-986187 binding site, and show how BMS-986187 binding leads to a conformational shift in the cholesterol binding pose. Orthosteric ligand, BMS-986187, cholesterol/lipid, receptor and key residues Y151 and W194 are colored in yellow, orange, forest green, palecyan, and magenta respectively.

Strikingly, we observe essentially the same state-dependent conformational changes of W194^4.50^ and Y151^3.34^ in all available high-resolution active and inactive structures of the closely related δ-opioid receptor (DOR)^20,34–39^ and κ-opioid receptor (KOR)^20,27,40–45^ (Extended Data Fig. 6A), suggesting that these may be conserved activation-associated rearrangements across opioid receptors. In nanobody-stabilized active-like states of both MOR and KOR, W194^4.50^ adopts a unique t-90 rotamer oriented toward the membrane (Extended Data Fig. 6B), while Y151^3.34^ displays receptor-specific rotameric conformations adopting a m-85 rotamer in MOR as in the respective inactive state, but a t-80 rotamer in KOR as observed in active states.

Overall, our analysis suggests that the binding pose of BMS-986187 observed in our DAMGO/MOR-G_i1_ complex cryo-EM structure is incompatible with the side chain orientations of Y151^3.34^ and W194^4.50^ seen in inactive (m-85 and m-95, respectively) and active-like (m-85 and t-90, respectively) states. While stabilization of the active state configuration may therefore contribute to BMS-986187 positive allosteric effects, it should be noted that lipids and sterols likely play an additional role. Indeed, one inactive MOR structure reveals a lipid occupying the allosteric site^24^, and previous MOR structures and our data show that cholesterol packs against W194^4.50^ in active G protein-bound states (Fig. 5B, Extended Data Fig 6C). This cholesterol molecule remains present in the BMS- 986187-bound DAMGO/MOR-G_i1_ complex structure, but shifts conformation, likely to accommodate the PAM (Extended Data Fig. 4A). While these observations offer initial insight into the structural basis of BMS-986187’s allosteric effects, the precise mechanisms by which it modulates orthosteric ligand potency, efficacy, or G protein engagement remain to be fully elucidated. This is underscored by the observation that the cholesterol binding pose is not conserved across opioid receptor subtypes (Extended Data Fig. 6C).

### A model of BMS-986187’s positive allosteric modulation of DAMGO-induced MOR signaling

To gain mechanistic insight into how BMS-986187 enhances orthosteric ligand activity and signal output at MOR, we investigated the allosteric communication pathway linking the PAM, the orthosteric ligand DAMGO, and the MOR-G_i1_ interface. Specifically, we analyzed our MD simulations of the DAMGO/MOR-G_i1_ signaling complex in the presence and absence of BMS-986187 using transfer entropy, an information-theory approach that quantifies the directionality of information flow between residue pairs over time, enabling inference of causality in residue dynamics^46^.

As summarized in Fig. 6, this analysis allowed us to trace how signaling propagates from BMS-986187, bound at the lipid-facing crevice between TM2, TM3, TM4 of MOR, to both the orthosteric site and the G protein-coupling interface. We observed three major signaling routes: (1) from BMS-986187 to DAMGO, potentially modulating DAMGO’s binding affinity (BMS-986187→DAMGO; Figs. 6B, 6E); (2) from DAMGO to the MOR–G_i1_ interface, potentially affecting efficacy (DAMGO→G protein; Figs. 6C, 6F); and (3) directly from BMS-986187 to the MOR–G_i1_ interface, potentially mediating PAM-driven agonism (BMS-986187→G protein; Figs. 6D, 6G).

**Fig 6.**
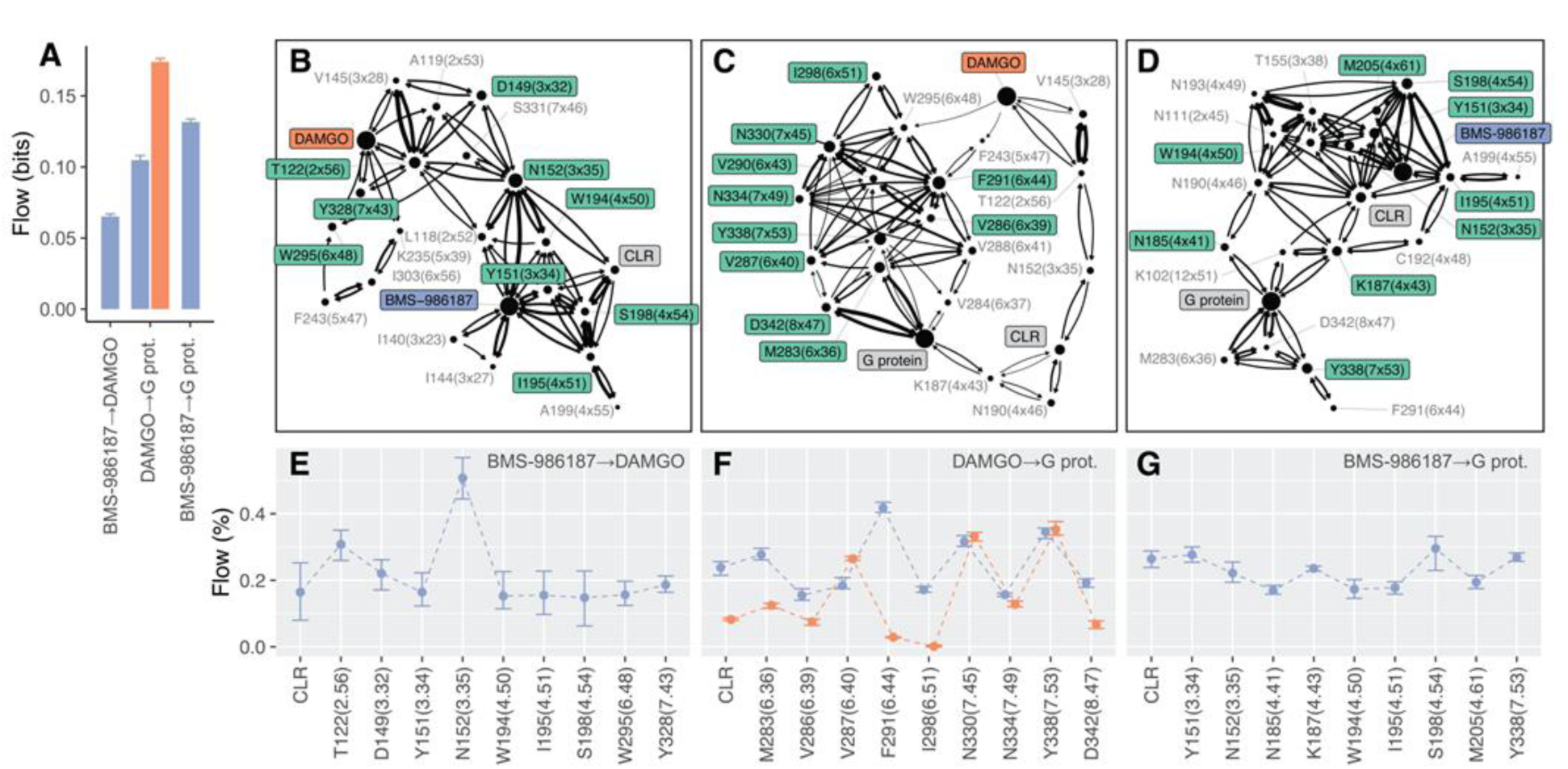
Pathway-specific information flow between BMS-986187, DAMGO and the G protein. (A) Total information flow (in bits) for three pathways: BMS-896187 to DAMGO, DAMGO to the G protein, and BMS-896187 to the G protein. (B–D) Network diagrams illustrating the predominant allosteric communication pathways within MOR: from BMS-986187 to DAMGO (B), from DAMGO to the G protein (C), and from BMS-896187 to the G protein (D). The 20 residues with the highest contribution to information flow are labeled, including the nearby cholesterol molecule (labeled CLR). The top 10 contributing residues are highlighted in green and enclosed in boxes. Node size is proportional to the total information flow transmitted or received by each residue, while edge thickness indicates the magnitude of transfer entropy between residue pairs. (E–G) Fractional information flow per residue (mean ± SEM, based on 20 samples of transfer entropy values) for the top 10 residues in each of the three pathways: BMS- 986187 to DAMGO (E), DAMGO to G protein (F), and BMS-986187 to G protein (G). In panel F, flow values in the absence of BMS-986187 are shown in orange for comparison.

Quantification of maximum information flux (Fig. 6A) revealed that BMS-986167 reduces the direct information flow from DAMGO to the G protein (compare purple with orange bars in Fig. 6A), possibly accelerating its communication pathway. Per-residue total flux analysis identified Y151^3.34^, W194^4.50^, and S198^4.54^, along with the cholesterol molecule adjacent to the PAM binding site, among the top 10 contributors to the BMS- 986187→DAMGO flux (Figs. 6B, 6E). Notably, Y151^3.34^, W194^4.50^, S198^4.54^, and the cholesterol molecule also ranked among the most prominent contributors to the BMS-986187*→G* protein communication pathway (Fig. 6D, 6G), further underscoring their central role in the PAM allosteric effects.

## Discussion

Allosteric modulators of opioid receptor function have emerged as promising tools to reshape opioid signaling modalities, offering pathways toward safer opioid-based treatments for pain, opioid overdose, and substance use disorders. While these compounds hold considerable therapeutic potential, only a limited number of allosteric modulators of opioid receptors have been identified to date^8–10^, and their underlying molecular mechanisms remain poorly understood.

Previous structural studies characterized the binding and mechanism of action of the MOR positive allosteric modulator BMS-986122^11^. Based on the absence of additive effects when BMS-986122 and BMS-986187 are co-applied, several groups hypothesized that the two PAMs act at a common site^12,13^. To test this hypothesis, we employed cryo-EM, MD simulations, and functional signaling assays to characterize the binding mode and mechanisms of BMS-986187. Unexpectedly, we found that BMS- 986187 binds to a membrane-facing crevice formed by TM2, TM3, and TM4, spatially distinct from the previously reported binding site of BMS-986122 (Fig. 2).

Notably, BMS-986187 also engages in interactions with a neighboring cholesterol molecule. Cholesterol is a well-known modulator of GPCR function^47^, has been implicated in modulating opioid receptor activity^48,49^, and prior studies have shown that cholesterol depletion can attenuate opioid receptor signaling^50^. Our cryo-EM structures and MD simulations consistently reveal a stably bound cholesterol molecule near the BMS- 986187 site, and importantly, show that binding of BMS-986187 alters the binding mode of cholesterol without displacing it. This raises important questions regarding the interplay between BMS-986187 and cholesterol, particularly in light of the W194^4.50^A mutation, which appears to potently enhance the PAM’s allosteric effects, while serving as a key binding site for cholesterol in the PAM-free state. It should be noted that interpreting the role of W194^4.50^ is further complicated by its dynamic behavior across MOR active and inactive states. In fact, our structural comparisons indicate that W194^4.50^ undergoes state- dependent rotameric changes, and such conformational changes may even be required for activation-related rotameric shifts in the neighboring Y151^3.34^ (Fig 5, Extended Data Fig 6). These observations highlight the complex relationship between BMS-986187 engagement, cholesterol modulation, and activation-related changes in MOR, and thus require a more detailed dissection of these mechanisms in future work.

It is also worth noting that BMS-986187 was originally reported to exhibit greater selectivity for DOR than for MOR (with additional allosteric activity at KOR)^13^. However, among the 10 MOR residues that make direct contact with BMS-986187, three (i.e., L118^2.52^, I144^3.27^, and L202^4.58^) are not conserved in DOR. While mutational studies and transfer entropy analysis suggest that these individual residues are not essential for BMS- 986187’s allosteric effects, their collective divergence may contribute to differences in PAM binding or signaling modulation at DOR. As a result, BMS-986187 may exert subtly distinct modulatory effects at DOR, a possibility that warrants further investigation. This distinction is particularly relevant given that cholesterol appears to play a more prominent role in MOR signaling than in DOR^50^. Notably, available DOR structures either lack a cholesterol molecule at the corresponding site^38,39^, or feature it in a noticeably different configuration (Extended Data Fig. 6C)^20^, which may in turn affect PAM binding and efficacy.

Despite these differences, DOR and even KOR structures display the same conserved rotamer changes as observed in MOR, irrespective of orthosteric ligand or structural method used. Of note, active-like MOR and KOR structures share a unique rotamer conformation of W^4.50^, providing further evidence that this residue may function as a structural gate during receptor activation.

In addition to W194^4.50^, our study implicates additional residues, including Y151^3.34^, M205^4.51^, and S198^4.54^, in receptor activation and/or in mediating the allosteric effects of BMS-986187 at MOR. However, it is important to recognize that our findings are based primarily on assays measuring MOR-mediated changes in cellular cAMP concentrations in response to DAMGO. Because GPCR signaling is highly pathway- and ligand-specific, the observed allosteric mechanisms uncovered here may vary depending on: i) the orthosteric agonist and/or ii) the signaling pathway, such as those stimulated by Gz, Go, arrestin, or other transducers. Our previous findings already indicated that MOR possess preference for distinct signal transducers^22^, and in related studies of the Gi/o/z-coupled 5-HT1A serotonin receptor we showed how distinct receptor conformations are responsible for engaging distinct G protein transducers^51^. The effects of BMS-986187 and other PAMs thus likely vary depending on receptor state and pathway. Furthermore, our previous work with MOR PAMs such as MS1 and compound 5 demonstrated substantial probe dependence, with allosteric effects varying strongly depending on the orthosteric agonist^8^. Similar findings were reported for BMS-986122, where differences were particularly pronounced between partial and full agonists^12^. Consistent with this, our transfer entropy analysis highlights directional information flow from BMS-986187 to DAMGO-specific interactions within the MOR binding site. These results suggest that i) BMS-986187 may differentially modulate the activity of distinct orthosteric opioids at MOR, and ii) modulators occupying distinct binding sites, such as BMS-986122, likely exhibit a distinct information flow signature. Future studies will be required to i) determine how BMS-986187 influences signaling elicited by other MOR agonists and across different pathways, and ii) define how these mechanisms diverge from those of BMS- 986122.

We hypothesize that such studies will uncover a unique information flow for each PAM that might in part explain why BMS-986187 and BMS-986122 do not display additive allosteric effects at MOR as previously hypothesized^12,13^. While such pharmacological non-additivity had been interpreted as evidence for overlapping binding sites, our structural data suggest that these chemically distinct PAMs likely act through orthogonal mechanisms. The absence of major conformational changes in the receptor across BMS- 986187-bound, BMS-986122-bound, and PAM-free MOR structures further implies that their effects are mediated by dynamic and pathway-specific mechanisms rather than large-scale structural rearrangements. Consequently, elucidating the divergent mechanisms of MOR PAMs will require an integrative approach that combines high-resolution structural biology, molecular dynamics simulations, and functional assays. Overall, our findings lay the groundwork for such future efforts and point toward the need for detailed mapping of receptor-, ligand-, and transducer-specific allosteric communication pathways. Ultimately, this work provides a framework for the rational design of allosteric modulators and integrative models capable of predicting how distinct ligands converge to shape GPCR signaling, an endeavor with far-reaching implications for the development of next-generation therapeutics.

## Methods

### Constructs and protein expression

For cryo-EM structure determination, a modified human MOR construct was cloned into pFastBac vector. We inserted a hemagglutinine signal sequence followed by a FLAG tag before the N-terminus of MOR, and a 3C protease cleave site followed by a 10 x Histidine tag was fused to the C-terminus. In addition, C-terminal residues (L389-P400) were removed due to flexibility and a thermostabilizing mutation F158^3.41^W was introduced^11^. For the G protein construct design, the heterotrimer was expressed from a single multibac virus. Human Gβ1 was cloned under the control of a polyhedrin promoter, and a Gγ2– Gαi1 fusion construct was cloned under the control of a P10 promoter, using a GSAGSAGSA linker to fuse Gγ2 and Gαi1. Four mutations (S47N, G203A, E245A and A326S) were introduced into Gαi1 to facilitate thermostability of the heterotrimeric Gi1 protein construct^16^.

MOR and G protein were co-expressed at an equal ratio in Sf9 cells (Expression Systems) using the Bac-to-Bac Baculovirus expression system. To obtain MOR and Gi1 viruses, the same amount of recombinant bacmid were incubated for 30 minutes with FuGENE HD transfection reagent (Promega) in SF900 II medium (Gibco). P0 was obtained by collecting supernatant 5 days after cells were infected with transfection mixture and P1 baculovirus was generated through transfecting cells with P0. Co-expression of MOR and Gi1 was done with P1 to infect cells at a density of 2 x 10^6^ per ml and a total multiplicity of infection of 5. The cells were cultured for two days at 27 °C and collected via centrifugation at 2,500 x g for 15 min, then stored at -80 °C until purification.

### Protein purification of complex of MOR and heterotrimeric Gi1

Frozen Sf9 cells were thawed on ice and disrupted by dounce homogenization in resuspension buffer containing 20 mM HEPES pH 7.5, 50 mM NaCl, 5 mM MgCl_2_, 5 mM CaCl_2_, 5% glycerol, 0.3 mM TCEP (Tris (2-Carboxyethyl) phosphine Hydrochloride, Gold Biotechnology) and home-made protease inhibitor cocktail (500 µM AEBSF, 1 µM E-64, 1 µM leupeptin and 150 nM aprotinin) (Gold Biotechnology). DAMGO was supplemented to a final concentration of 20 µM, and BMS-986187 was added at 3 µM for the PAM- bound sample. Cell membrane suspensions were incubated with ligands for 30 minutes to facilitate the formation of MOR-Gi1 complexes. Apyrase at a working concentration of 25 mU/mL for another 30 minutes was used to digest nucleotides and prevent complex disassembly. Cell membranes were spun down at 40,000 rpm for 25 minutes after incubation.

Protein complexes were extracted in solubilization buffer containing 20 mM HEPES pH 7.5, 50 mM NaCl, 5 mM MgCl_2_, 0.3 mM TCEP, 1% (w/v) n-dodecyl-β-d- maltopyranoside (DDM; Anatrace), 0.2% (w/v) cholesteryl hemisuccinate (CHS; Anatrace) and home-made protease inhibitor cocktail. Protein samples were subsequently collected by centrifugation at 55,000 rpm for 25 min after solubilization. TALON IMAC resin (Clontech) and 20 mM imidazole (Sigma) were applied to supernatant for overnight incubation at 4 °C.

The following day, protein-bound resin was washed with 15 column volumes of wash buffer I (20 mM HEPES, pH 7.5, 50 mM NaCl, 5 mM MgCl_2_, 0.1% (w/v) DDM, 0.02% (w/v) CHS, 20 mM imidazole, 5% (v/v) glycerol, 0.3 µM TCEP and 10 µM DAMGO). 3 µM BMS- 986187 was separately added in all purification buffer for Pam-bound sample. Then detergent was exchanged to lauryl maltose neopentyl glycol (LMNG) with wash buffer I supplemented with 0.1% LMNG. The protein-bound resin was equilibrated with 10 column volumes of LMNG buffer (20 mM HEPES, pH 7.5, 50 mM NaCl, 0.5% (w/v) LMNG, 0.002% (w/v) CHS, 20 mM imidazole, 5% (v/v) glycerol, 0.3 µM TCEP and 10 µM DAMGO) followed by a washing step with 10 column volumes of wash buffer II (20 mM HEPES, pH 7.5, 50 mM NaCl, 0.5% (w/v) LMNG, 0.002% (w/v) CHS, 20 mM imidazole, 5% (v/v) glycerol, 0.3 µM TCEP and 10 µM DAMGO), and subsequent additional 10 column volumes of wash buffer III (10 mM HEPES, pH 7.5, 30 mM NaCl, 0.01% (w/v) LMNG, 0.002% (w/v) CHS, 20 mM imidazole, 5% (v/v) glycerol, 0.3 µM TCEP and 20 µM DAMGO). The protein samples were then gradually eluted in fractions using 1 column volume of elution buffer (10 mM HEPES, pH 7.5, 30 mM NaCl, 0.01% (w/v) LMNG, 0.002% (w/v) CHS, 250 mM imidazole, 5% (v/v) glycerol, 0.3 µM TCEP and 20 µM DAMGO). The best fractions were selected according to an initial assessment on size exclusion chromatography and SDS-PAGE. The selected fractions were concentrated to ∼500 µL volume with Vivaspin 6 centrifugal concentrators (Sartorius) and applied to PD MiniTrap desalt column (Cytiva). The desalted protein samples were concentrated to below 100 µL and further purified over a Superdex 200 Increase size exclusion column (Cytiva) equilibrated in 20 mM HEPES, pH 7.5, 50 mM NaCl, 0.001%(w/v) LMNG, 0.0002% (w/v) CHS, 0.0002% GDN, and 20 µM DAMGO. Peak fractions were pooled, concentrated to ∼1.5-3 mg/ml, and immediately used to prepare grids for cryo-EM data collection.

### Cryo-EM sample preparation for MOR-Gi1 complexes

3 µL of the purified protein complex, both with and without BMS-986187, were applied onto glow-discharged holey carbon EM grids (Quantifoil 300 copper mesh, R1.2/1.3) in an EM-GP2 plunge freezer (Leica). Sample freezing was performed at 95% humidity and 4 °C with a blotting force setting to 0. The grids were stored in liquid nitrogen for cryo-EM data collection and analysis after plunge-freezing into liquid ethane.

### Cryo-EM data collection

All automatic data collection was performed on a FEI Titan Krios instrument equipped with a Gatan K3 direct electron detector operated by the Simons Electron Microscopy Center at the New York Structural Biology Center (New York, New York). For the DAMGO/MOR-Gi1 complex structure, the microscope was operated at 300 keV accelerating voltage, at a nominal magnification of 64,000x corresponding to a pixel size of 1.076 Å. 8,259 movies were obtained at a dose rate of 25.97 electrons per Å2 per second with a defocus ranging from −0.8 to −2.0 μm. The total exposure time was 2 s and intermediate frames were recorded in 0.05 s intervals, resulting in an accumulated dose of 51.94 electrons per Å2 and a total of 40 frames per micrograph. For the DAMGO/MOR-Gi1 complex bound to BMS-986187, the microscopes were operated at 300 keV accelerating voltage, at a nominal magnification of 105,000 corresponding to pixel sizes of 0.844 Å. 12,785 movies were obtained at a dose rate of 33.82 electrons per Å2 per second with a defocus ranging from −0.7 to −2.2 µm. The total exposure time was 1.5 seconds, and intermediate frames were recorded in 0.05 s intervals, resulting in an accumulated dose of 50.74 electrons per Å2 and a total of 30 frames per micrograph.

### Cryo-EM data processing and model refinement

For each dataset, movies were motion-corrected using MotionCor2^52^ and imported to cryoSPARC^53^ for further processing. Contrast transfer functions were estimated using patchCTF in cryoSPARC. An initial map template was produced from a subset of micrographs using blob picking, followed by particle extraction, 2D classification and generation of a model ab initio. We observe that our DAMGO/MOR-Gi1 complexes cryo-EM form antiparallel dimers as have been observed before in previous opioid receptor cryo-EM studies^14,20^. Following template generation, subsequent map models were produced from a curated micrograph set using particles found by picking using the initial map template. Bad models were generated from rejected particles to remove damaged particles or debris from good particles during several successive 3D classification and heterorefinement jobs. For the BMS-986122-bound DAMGO/MOR-Gi1 complex we obtained a final particle stack of 101,558 particles. We next performed a final non-uniform (NU) refinement^54^ step using C2 symmetry of the antiparallel dimer, resulting in a map with a final global resolution of 3.47 Å. Finally, we further used the DeepEMhancer^55^ tool in cryoSPARC to generate a final map, which resulted in an overall improvement of the map, but lowered the quality of the density for some moieties of BMS-986187 (Extended Data Fig 2). We did attempt to further improve the maps by making masks to remove the G proteins and performing local refinement on only the receptor dimer, or just one of the receptor protomers. This, however, did not improve the map quality of the receptor or BMS-986187, and so we opted to use the initial maps using only the standard dynamic mask from NU-refinement that did include the G proteins. A similar processing strategy for the PAM-free DAMGO/MOR-Gi1 complex resulted in a final map containing 791,402 particles, with a final global resolution of 2.89 Å.

Structures for both complexes were subsequently built in Coot^33^ starting with a model based on the BMS-986122-bound complex structure (PDB 8K9L)^11^. The starting model was docked into the corresponding cryo-EM maps using UCSF ChimeraX^56^, and the models were manually adjusted in Coot and further refined in PHENIX^57^. A cif file and starting model for BMS-986187 was generated using Grade from global phasing (https://grade.globalphasing.org). A final model of BMS-986187 bound to the DAMGO/MOR-Gi1 complex was then built based on the two maps of this dataset outlined above. Overall, we observe considerably lower map quality for the G proteins compared to the receptor and BMS-986187-binding site. However the density was sufficient to place the backbone of Gβ1, Gγ2, and most of Gαi1 at lower contour levels, and we therefore only omitted the Gαi1 alpha helical domain (AHD). Since we do not analyze details of the G protein as part of this study, we felt that our data was sufficient to warrant modeling of the entire heterotrimer except for the Gαi1 AHD.

### cAMP inhibition assay

Gi/o-mediated cAMP reduction was measured in HEK293-T cells (American Type Culture Collection) using a cAMP biosensor^58^ (GloSensor, Promega). HEK293-T cells were cultured before transfection in Dulbecco’s modified Eagle medium (DMEM) supplemented with 10% (v/v) FBS and penicillin–streptomycin (Invitrogen). Wild type MOR or mutants along with the cAMP biosensor were transfected into cells in a 10 cm plate when cells reached ∼70% confluence. 0.8 μg receptor DNA and 8 μg GloSensor DNA formed particles with 16 μL of polyethyleneimine (PEI; Alfa Aesar) in 500 µl Opti-MEM (Invitrogen) and were added directly onto cells. The following day, cells were resuspended and plated in 384-well plates (Greiner Bio One) at 20,000 cells per well in DMEM supplemented with 1% (v/v) dialyzed FBS (dFBS) and penicillin–streptomycin. After 24h, media was removed and 0.4 mg/ml D-luciferin (Gold Biotechnology) was loaded into cells in 15 µl of Hank’s balanced salt solution (HBSS) supplemented with 20 mM HEPES pH 7.4, 0.1% BSA (w/v) and 0.01% ascorbic acid (w/v), and cells were incubated for 1 hr. DAMGO and BMS-986187 were prepared separately at 3x concentration in HBSS supplemented with 20 mM HEPES pH 7.4, 0.1% BSA (w/v) and 0.01% ascorbic acid (w/v), and 15 µl of each were added to the cells and incubated for 30 minutes in the dark. Forskolin (MedChem) was then added to a final concentration of 1 nM to elevate cellular cAMP levels, and luminescence intensity was determined 45 min following forskolin addition on a MicroBeta TriLux liquid scintillation counter (Perkin Elmer). Experiments were performed in triplicate and performed independently at least three times. Data were plotted and analyzed using GraphPad Prism 8.0.

### BRET signaling assay

For transfection, HEK293-T cells were transfected with wild type MOR and heterotrimeric Gi1 (Gαi1/Gβ3/Gγ9) TRUPATH reporter construct^59^ at the ration of 1:3:3:3, using 0.8 μg receptor DNA and 2.4 μg of each G protein. The next day, cells were transferred into a 384-well plate at 20,000 cells per well. The following day cells were washed in HBSS supplemented with 20 mM HEPES pH 7.4, 0.1% BSA (w/v) and 0.01% ascorbic acid (w/v), and stimulated with different drug concentrations. Cells were incubated with drugs at 37 °C for 30 minutes followed by the addition of coelenterazine 400a, which was prepared freshly and used at the final concentration of 5 μM. Plates were then immediately read in a Victor NIVO plate reader (Perkin Elmer) with 395 nm (RLuc8- coelenterazine 400a) and 510 nm (GFP2) emission filters, at integration times of 1 s per well. Net BRET signals were computed as the ratio of the GFP2 emission to RLuc8 emission and analyzed in GraphPad Prism 8.0. Experiments were performed in triplicate and performed independently at least three times.

### System setup for MD simulations

MD simulations were initiated using nucleotide-free, DAMGO-bound MOR-G_i1_ complex structures that had been preliminarily fitted to cryo-EM density maps prior to final refinement. These structures included or excluded the positive allosteric modulator BMS- 986187 and contained a nearby cholesterol molecule. The missing Gα_i1_ AHD (residues 56–181), unresolved in the cryo-EM map, was modeled in an open conformation using the neurotensin receptor 1–G_i1_ structure (PDB ID: 7L0Q)^60^ as a template, after aligning its Gα_i1_ Ras-like helical domain (RHD) with the corresponding region in the DAMGO/MOR–G_i1_ complexes.

Unresolved segments, including MOR helix 8 (residues 343–347), Gα_i1_ N-terminus (residues 1–4), and Gγ C-terminus (residues 64–68), were modeled either by homology using available high-resolution crystal structures (e.g., PDB ID: 1GP2^61^ for Gα_i1_ and 4DKL^24^ for MOR) or built de novo using MODELLER^62^.

System preparation was performed using CHARMM-GUI^63^ with the CHARMM36m force field^64^, employing hydrogen mass repartitioning to enable an extended integration timestep^65^ Residue D114^2.50^ was protonated, while all other titratable residues were assigned protonation states corresponding to pH 7.

To mimic physiological membrane anchoring, lipid modifications were included as previously described^66^: Gα_i1_ was myristoylated at G2 and palmitoylated at C3, and Gγ was geranylgeranylated at C68. The protein complexes were embedded in a mixed lipid bilayer designed to reflect the composition of a mammalian plasma membrane^67^. This bilayer included 1-palmitoyl-2-oleoyl-sn-glycero-3-phosphocholine (POPC), 1-palmitoyl- 2-oleoyl-sn-glycero-3-phosphoethanolamine (POPE), 1-palmitoyl-2-oleoyl-sn-glycero-3- phosphoserine (POPS), palmitoyl sphingomyelin (PSM), monosialodihexosylganglioside (GM3), cholesterol, and 1,2-diacyl-sn-glycero-3-phospho-1-D-myo-inositol 4,5- bisphosphate (PIP2), all known to influence GPCR function and G protein coupling ^67^ Systems were solvated with TIP3P water molecules and neutralized with 0.15 M NaCl to emulate physiological ionic strength.

### MD simulation protocol and analyses

All ligands were parameterized with the CHARMM General Force Field and MD simulations were carried out using Gromacs 2023^68^. Each system was initially subjected to 5000 steps of steepest descent energy minimization, followed by a multi-phase restrained equilibration protocol. The first 5 ns of equilibration involved gradually reducing positional restraints (starting at 1000 kJ/mol) on all non-solvent, non-hydrogen atoms while maintaining full restraints on protein and ligand heavy atoms. This was followed by a 200 ns equilibration phase with restraints applied only to protein and ligand heavy atoms to allow proper relaxation of the lipid bilayer.

Production simulations consisted of 20 independent replicas per system, each run for 500 ns. The initial 50 ns of each trajectory were discarded, and snapshots were saved every 1 ns for downstream analysis.

Trajectory analyses and residue dynamics featurization for transfer entropy estimation were conducted using the Python libraries mdtraj (1.10.3)^69^ and scikit-learn (1.6.1)^70^. Ligand-protein and protein-protein interactions were calculated for each frame using ProLIF^71^ and were assigned either to the protein backbone or to the sidechain for protein residues. Interacting ligand atoms were grouped by fragments calculated using the BRICS algorithm^72^ implemented in RDKit^73^. Within each snapshot, duplicate interactions were counted only once, and only interactions present in >35% of the analyzed frames were reported. Confidence intervals were estimated by bootstrapping averages from 25 ns trajectory blocks.

To improve computational efficiency in featurizing residue dynamics and estimating transfer entropy, we initially filtered residue pairs by retaining only those with minimum heavy-atom distances below 15 Å in the initial structure of each trajectory. Each residue’s conformation was featurized based on binary contact patterns (minimum heavy-atom distance < 4.5 Å) with other residues. We then applied k-means clustering (10 clusters per residue, initialized independently) to define a discrete conformational state variable 𝑋_𝑡,𝑖_ for each residue 𝑖 at time 𝑡. The orthosteric ligand DAMGO, the allosteric modulator BMS-986187, and the cryo-EM-resolved cholesterol molecule adjacent to the PAM site were included in the analysis and treated as protein residues.

To quantify the information flow between residues 𝑖 and 𝑗, we computed the transfer entropy as a Kullback-Leibler divergence ^46^.

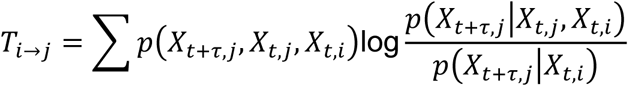

using a time lag 𝜏 = 1 ns (equal to the trajectory saving frequency). To correct for small-sample bias, we computed effective transfer entropy^74^ defined as 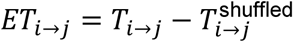 where 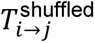 is calculated using a time-shuffled version of the series for process 𝑗, eliminating both temporal and statistical dependencies. The shuffled estimates were averaged across multiple iterations to robustly estimate and subtract sampling bias.

To assess statistical significance of transfer entropy values, we applied a Markov block bootstrap approach, as described by Dimpfl and Peter (2013)^75^. Transfer entropy computations were performed using the RTransferEntropy (0.2.21) package^76^, and network analyses were conducted with igraph (1.4.2^77^). Once all pairwise effective transfer entropy values were computed, we filtered out those with p-values greater than 10^-6^, retaining only statistically significant edges. To build the information flow network, we included only edges between residues in direct contact (≤ 4.5 Å) and used effective transfer entropy values as edge capacities.

We then computed the maximum information flux^78^ from BMS-9862187 to the DAMGO site, and from both ligands to G protein residues at the receptor interface. For each information flux calculation, we also quantified the total flux through each residue, defined as the sum of incoming and outgoing fluxes.

## Acknowledgements

This work was supported by NIH grants R01DA058681 (D.W.) and R01DA063209 (M.F.), an Irma T. Hirschl/Monique Weill-Caulier Trust Research Award (D.W.), NIH F31 fellowship MH132317 and T32 Training Grant GM062754 (A.L.W), and T32 Training Grant DA053558 (G.Z.). Some of this work was performed at the National Center for cryo-EM Access and Training (NCCAT) and the Simons Electron Microscopy Center located at the New York Structural Biology Center, supported by the NIH Common Fund Transformative High Resolution Cryo-Electron Microscopy program (U24 GM129539,) and by grants from the Simons Foundation (SF349247) and NY State Assembly. We further acknowledge cryo-EM resources at the National Resource for Automated Molecular Microscopy located at the New York Structural Biology Center, supported by grants from the Simons Foundation (SF349247), NYSTAR, and the NIH National Institute of General Medical Sciences (GM103310) with additional support from Agouron Institute (F00316) and NIH (OD019994). Computational work was supported in part through the computational resources and staff expertise provided by Scientific Computing at the Icahn School of Medicine at Mount Sinai and supported by the Clinical and Translational Science Awards (CTSA) grant UL1TR004419 from the National Center for Advancing Translational Sciences. Research reported in this paper was supported by the Office of Research Infrastructure of the National Institutes of Health under award number S10OD026880 and S10OD030463. The content is solely the responsibility of the authors and does not necessarily represent the official views of the National Institutes of Health. We would also like to acknowledge J. McCorvy for critical evaluation of the manuscript.

## Author Contributions

H.Z. designed experiments, collected data, refined structures, performed signaling assays, and wrote the manuscript. K.K. performed computational simulations and analyzed them with help from D.P. A.K.P. prepared samples for cryo-EM with help from G.Z. and A.L.W. S.Y. and S.W. helped with signaling assays. A.A. helped with data refinement. M.F. designed and supervised computational experiments, analyzed related data, and wrote the manuscript. D.W. designed experiments, analyzed the data, supervised the project, and wrote the manuscript.

## Data Availability

Electrostatic potential maps and structure coordinates have been deposited in the Electron Microscopy Data Bank (EMDB) and the Protein Data Bank (PDB) under accession codes EMD-71871/PDB-9PUD (DAMGO/MOR-Gαi1-Gβ1-Gγ2) and EMD- 71869/PDB-9PU5 (BMS986187/DAMGO/MOR-Gαi1-Gβ1-Gγ2).

## Extended Data

**Extended Data Fig 1.**
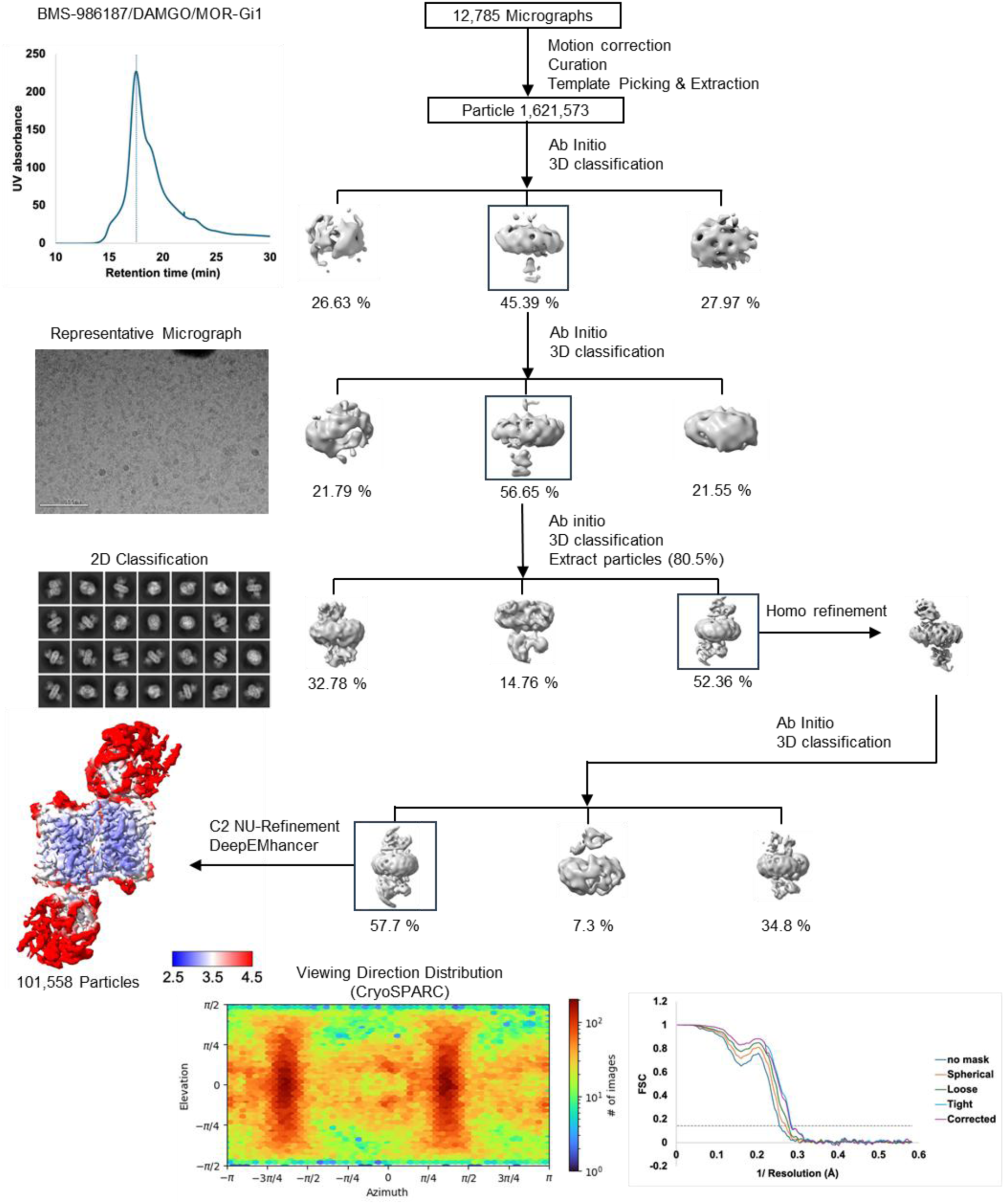
Purification, and cryo-EM structure determination of the BMS-986187-bound DAMGO/MOR-Gi1 complex. Analytical size exclusion chromatogram shows largely monodisperse BMS-986187/DAMGO/MOR-Gi1 complex sample. Data were collected on a 300 keV Krios, a representative micrograph is shown, and processed in cryoSPARC v4.1.2.: Particles were picked from motion corrected micrographs, subjected to 2D classification (representative classes are shown), followed by ab initio model building and 3D classification. After multiple rounds of 3D classification, the final particle stack of 101,558 particles was subjected to non-uniform refinement applying C2 symmetry. A final map was obtained using DeepEMhancer and GS-FSC indicates a global resolution of 3.47 Å applying the 0.143 cutoff. Viewing direction distribution analysis (cryoSPARC) indicates sufficient coverage. Calculations in cryoSPARC indicate local resolutions of up to 3 Å around the BMS-986187 binding site.

**Extended Data Fig 2.**
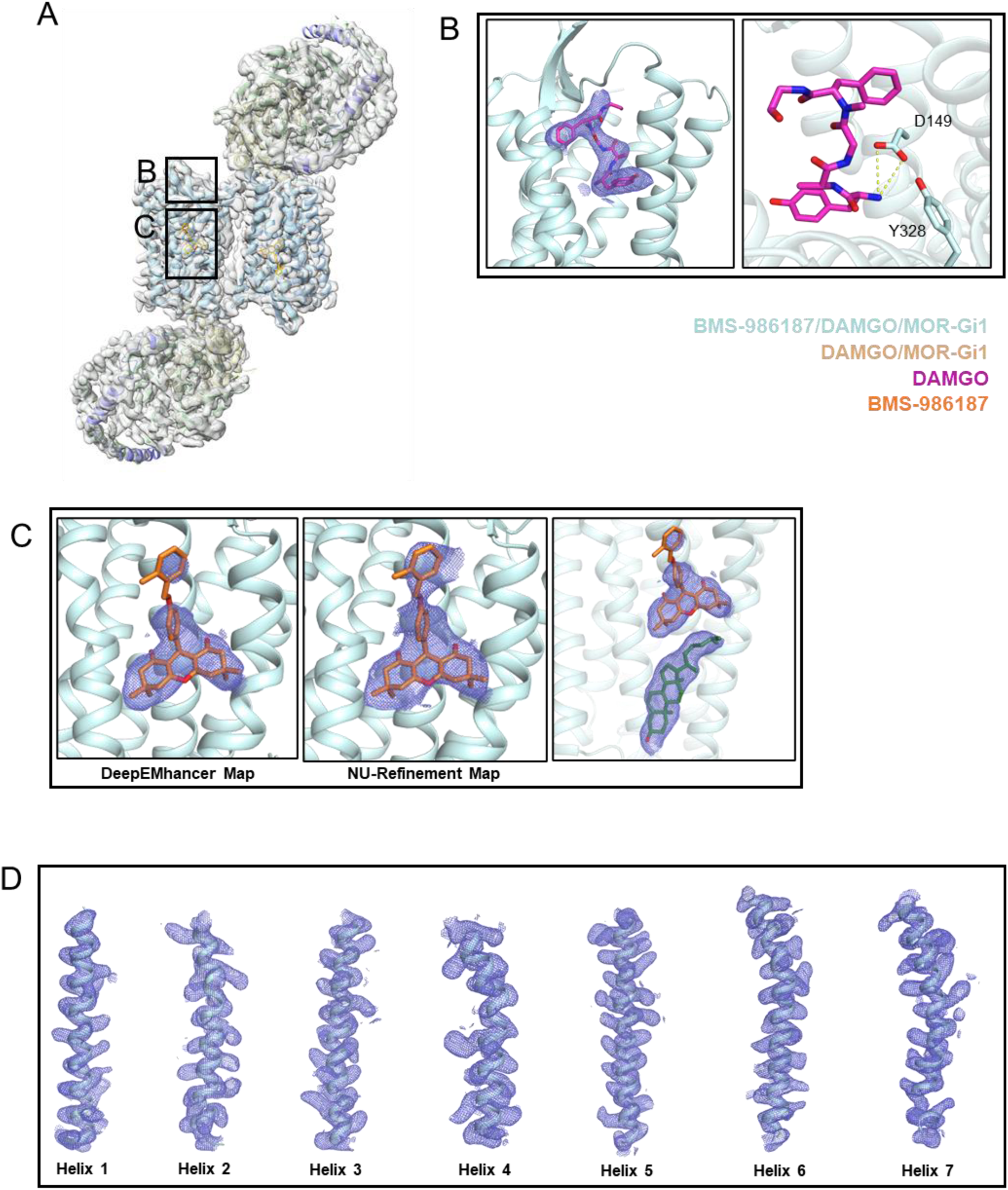
Representative map density of BMS-986187/DAMGO/MOR-Gi1 complex cryo-EM structure. (A) Overall cryo-EM map processed using DeepEMhancer and fitted with BMS-986187-bound DAMGO/MOR-Gi1 model built in COOT and refined in PHENIX. Inlays highlight map densities for DAMGO (B), BMS-986187 and cholesterol (C), and representative areas of MOR (D). MOR, DAMGO, BMS-986187, and cholesterol are shown in palecyan, magenta, orange, and green, respectively. Densities are shown as blue mesh with contour levels of 4.0σ for DAMGO (B), 3.0σ for the DeepEMhancer map, 3.5σ for the NU-refinement map, and 3.0σ for the maps of cholesterol and BMS-986187 (C), as well as 3.4σ for representative areas of MOR (D).

**Extended Data Fig 3.**
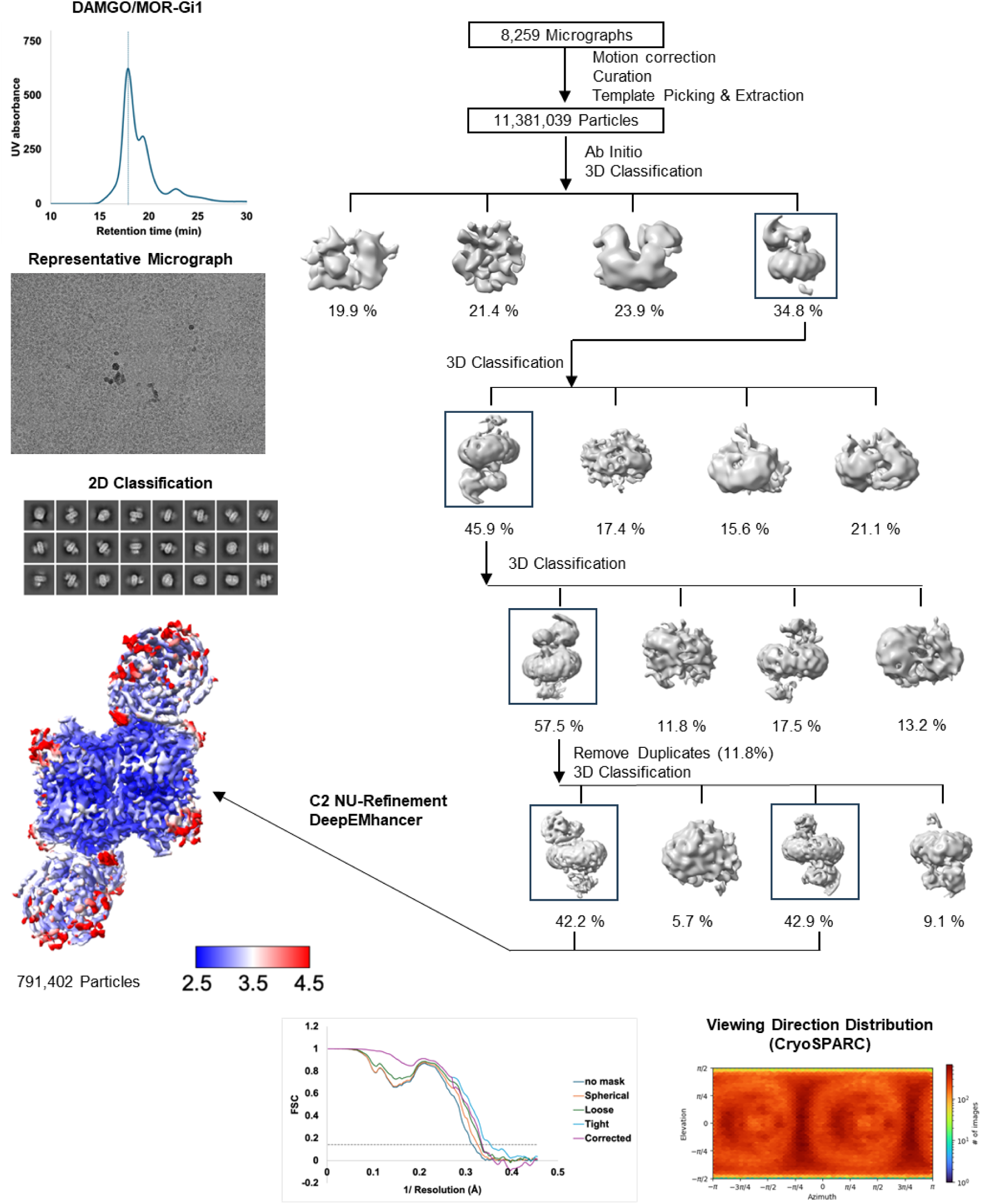
Purification, and cryo-EM structure determination of the PAM-free DAMGO/MOR-Gi1 complex. Analytical size exclusion chromatogram shows largely monodisperse DAMGO/MOR-Gi1 complex sample. Data were collected on a 300 keV Krios, a representative micrograph is shown, and processed in cryoSPARC v4.1.2.: Particles were picked from motion corrected micrographs, subjected to 2D classification (representative classes are shown), followed by ab initio model building and 3D classification. After multiple rounds of 3D classification, the final particle stack of 791,402 particles was subjected to non-uniform refinement applying C2 symmetry. A final map was obtained using DeepEMhancer and GS-FSC indicates a global resolution of 2.89 Å applying the 0.143 cutoff. Viewing direction distribution analysis (cryoSPARC) indicates sufficient coverage. Calculations in cryoSPARC indicate local resolutions of up to 2.5 Å around the BMS-986187 binding site.

**Extended Data Fig 4.**
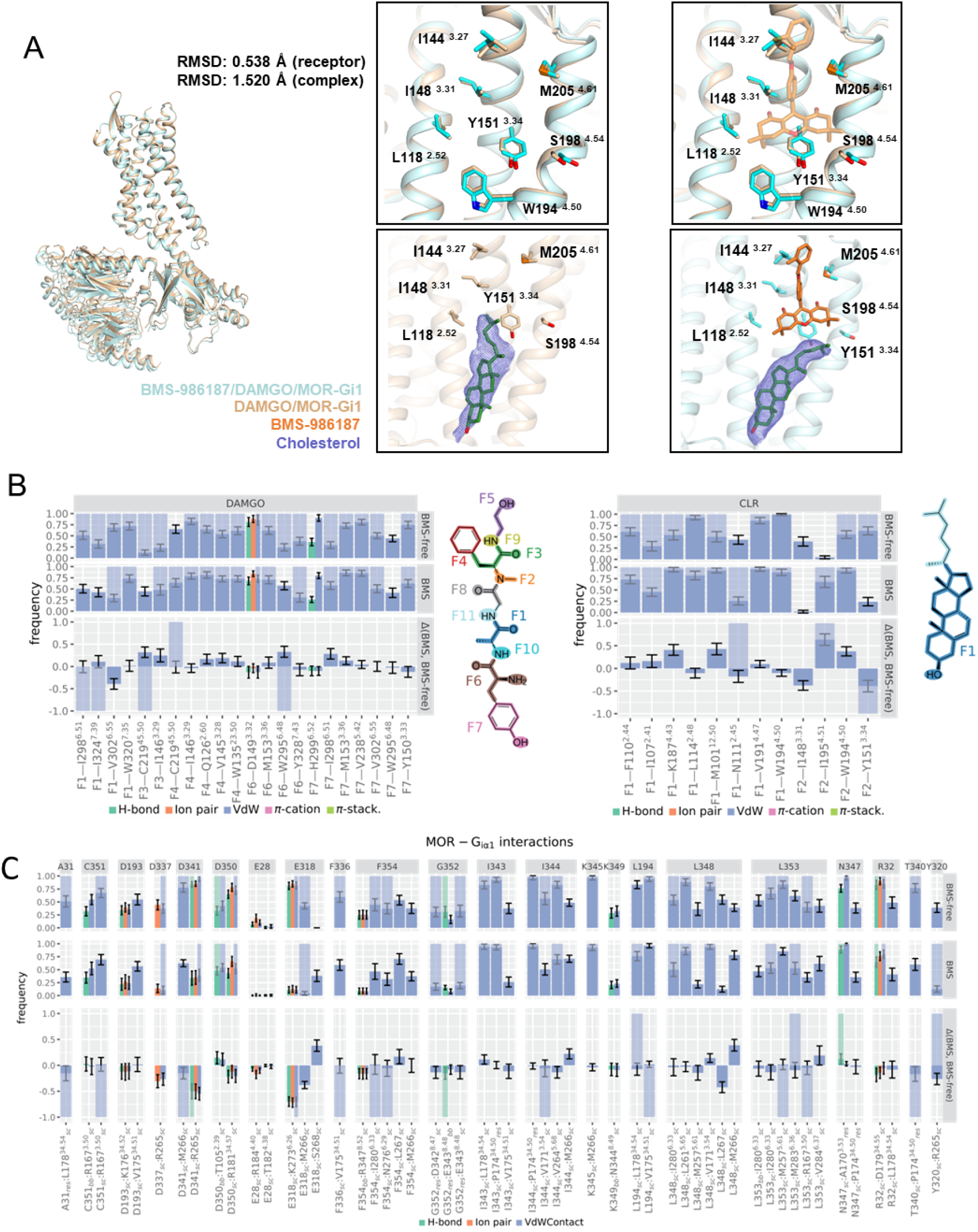
Effect of BMS-986187 on interactions within the MOR signaling complex. (A) Comparison of BMS-986187-bound (palecyan) and PAM-free (wheat) DAMGO/MOR-Gi1 cryo-EM structures. Top panel shows zoom in of BMS-986187 binding site and key residues, and bottom panel highlights conformational shift of cholesterol upon BMS-986187 binding. Densities for cholesterol are shown as blue mesh with a contour level of 4.0σ (PAM-free state) and 3.0σ (BMS-986187-bound state). (B–D) Comparison of interaction frequencies between DAMGO and MOR as well as cholesterol (CLR) and MOR (B), and interaction frequencies between Gαi1 and MOR (C) in molecular dynamics (MD) simulations with and without BMS-986187. Bar height represents the average interaction frequency across the trajectory, with error bars indicating the standard deviation calculated over 25 ns time blocks. Blue shading over a bar denotes interactions also observed in the cryo-EM structure. Each panel contains three facets: the top two show interaction frequencies in BMS-free and BMS-bound simulations, respectively; the bottom facet shows the difference in interaction frequency between the two conditions. Molecular fragmentations of DAMGO and CLR used for grouping interaction data are shown in B.

**Extended Data Fig 5.**
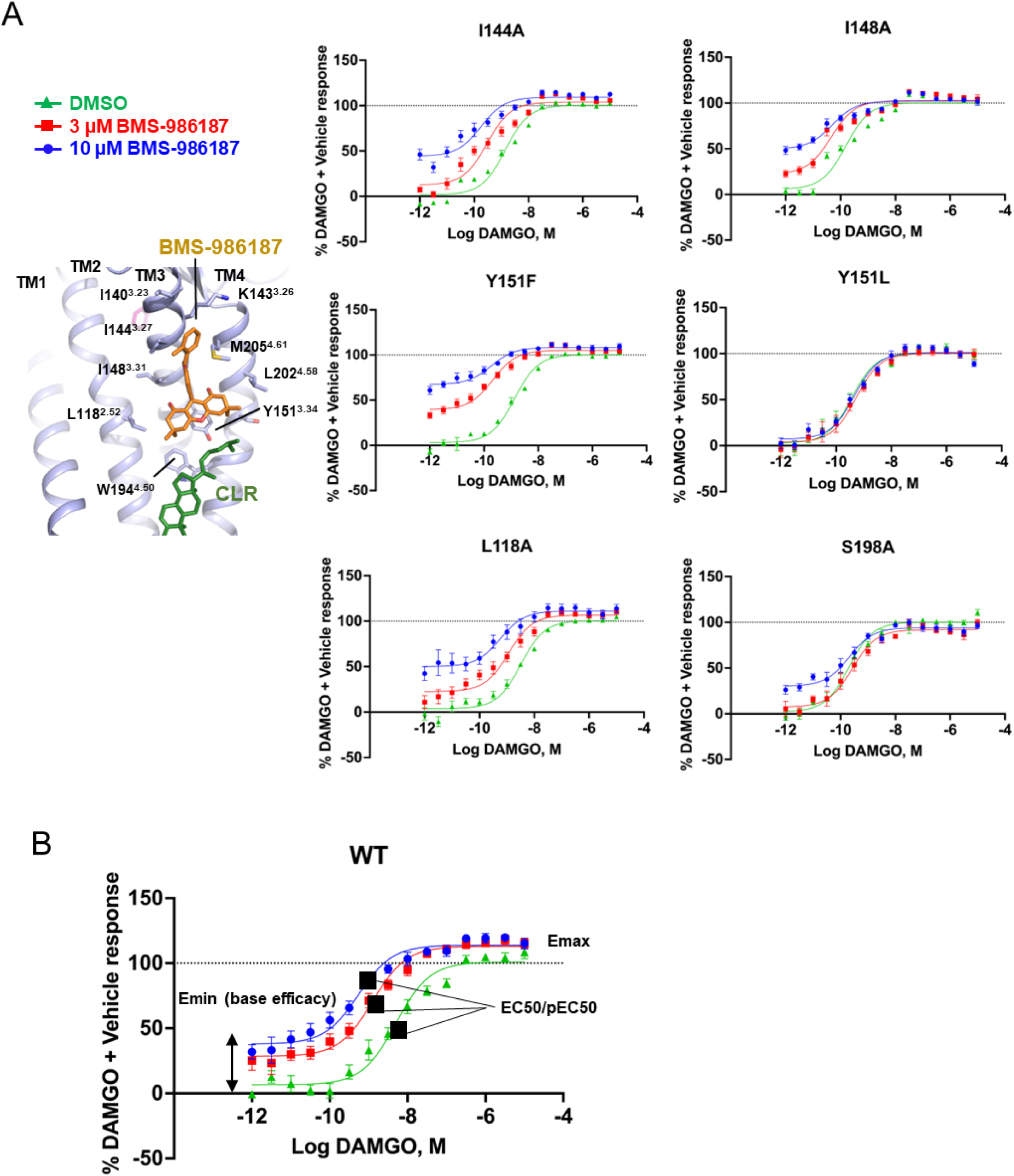
Effect of site directed receptor mutations on BMS-986187-modulated DAMGO activity at MOR as measured by cAMP accumulation. (A) Zoom in on BMS-986187 and key residues in the PAM binding site and DAMGO concentration response data in the presence of vehicle (DMSO, green), 3 µM BMS-986187 (red), and 10 µM BMS-986187 (green). All data from signaling studies represent mean ± SEM of three to five independent experiments (n = 3-5) performed in triplicate, and have been normalized to the DAMGO + vehicle response. (B) Illustration of DAMGO potency and efficacy. BMS-986187-mediated effects are measured by increases in DAMGO potency (pEC50) and base efficacy (Emin).

**Extended Data Fig 6.**
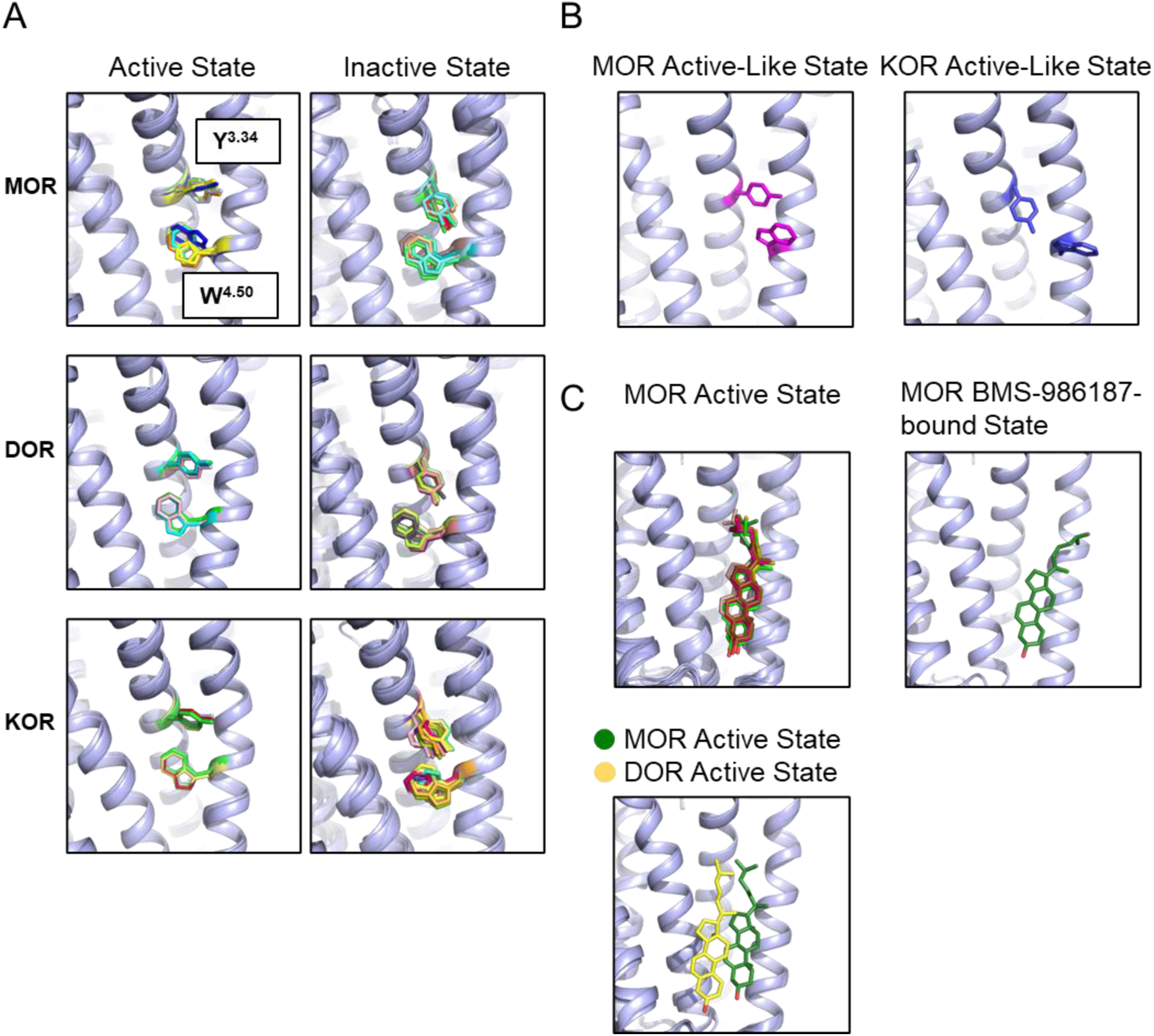
Structural analysis of Y^3.34^ and W^4.50^ conformations across different MOR, DOR, and KOR activation states. (A) Structures of inactive- and active-state opioid receptor structures (light blue) shown in the left and right columns respectively. Y^3.34^ and W^4.50^ are highlighted as sticks, and have been colored by structure (*<u>MOR Inactive State Structures</u>* – PDB: 4DKL (light blue), PDB: 7UL4 (limon), PDB: 8QOT (pale yellow), PDB: 9BJK (dark blue), PDB: 9MQH (yellow), PDB: 9MQI (orange), PDB: 9MDJ (cyan); *<u>MOR Active State Structures</u>* – PDB: 6DDE (green), PDB: 6DDF (red), PDB: 7SBF (wheat), PDB: 7SCG (light teal), PDB: 7T2G (dark salmon), PDB: 7T2H (pale green), PDB: 7U2K (light orange), PDB: 7U2L (purple), PDB: 8EF5 (brown), PDB: 8EF6 (forest green), PDB: 8EFB (dark red), PDB: 8EFL (pink), PDB: 8EFO (olive), PDB: 8EFQ (pale cyan), PDB: 8F7R (light pink), PDB: 8K9L (green cyan); *<u>DOR Inactive State Structures</u>*– PDB: 4EJ4 (aquamarine); PDB: 4N6H (green); PDB: 4RWA (light salmon); PDB: 4WRD (marine); *<u>DOR Active State Structures</u>* – PDB: 6PT2 (mint green), PDB: 6PT3 (light orange), PDB: 8F7S (salmon), PDB: 8Y45 (grey), PDB: 9CGJ (violet), PDB: 9CGK (limon); *<u>KOR Inactive State Structures</u>* – PDB: 4DJH (light blue), PDB: 6VI4 (green), PDB: 9MQK (yellow orange), PDB: 9MQL (dark red); *<u>KOR Active State Structures</u>* – PDB: 7YIT (pink), PDB: 8DZP (pale yellow), PDB: 8DZQ (bright orange), PDB: 8DZR (chartreuse), PDB: 8DZS (salmon), PDB: 8F7W (light orange), PDB: 8VVE (violet), PDB: 8VVF (light salmon), PDB: 8VVG (lime green), PDB: 9D61 (brown)). (B) MOR (PDB: 5C1M) and KOR (PDB: 6B73) nanobody-stabilized active-like state structures, with key residues colored in magenta and purple, respectively. (C) Cholesterol binding poses in MOR active state structures (PDB: 7T2G (dark salmon), PDB: 8EF5 (brown), PDB: 8EF6 (green), PDB: 8EFL (pink), PDB: 8EFO (olive), PDB: 9PUD (forest green)) and the BMS-986187-bound MOR active state (PDB: 9PU5 (forest green)), in the top left and right panels respectively. Comparison of cholesterol binding poses in active state structures of DOR (PDB: 8F7S (yellow)) and MOR (PDB: 5PUD (forest green)) shown in the bottom left panel.

**Extended Data Table 1.** Cryo-EM data collection, refinement and validation statistics.

|  | **BMS986187/**<br>**hMOR-Gi1-DAMGO**<br>**(EMD-71869)**<br>**(PDB ID: 9PU5)** | **hMOR-Gi1-DAMGO**<br>**(EMD-71871)**<br>**(PDB ID: 9PUD)** |
| --- | --- | --- |
| **Data collection and processing** |  |  |
| Magnification | 105,000 | 64,000 |
| Voltage (kV) | 300 | 300 |
| Electron exposure (e-/Å<sup>2</sup>) | 50.74 | 51.94 |
| Defocus range (µm) | -0.7 ~ -2.2 | -0.8 ~ -2.0 |
| Pixel size (Å) | 0.844 | 1.076 |
| Symmetry imposed | C2 | C2 |
| Initial particle images (no.) | 7,018,231 | 12,494,868 |
| Final particle images (no.) | 101,558 | 791,402 |
| Map resolution (Å) | 3.47 | 2.89 |
| FSC threshold | 0.143 | 0.143 |
| Map sharpening B-factor (Å<sup>2</sup>) | -152.8 | -140.9 |
| **Refinement** |  |  |
| Model composition |  |  |
| Non-hydrogen atoms | 14466 | 14284 |
| Protein residues | 1808 | 1808 |
| Ligands | 2 | 1 |
| R.m.s. deviations |  |  |
| Bond lengths (Å) | 0.003 | 0.006 |
| Bond angles (°) | 0.531 | 0.766 |
| Validation |  |  |
| Clashscore | 7.89 | 14.73 |
| Poor rotamers (%) | 0 | 0 |
| Ramachandran plot |  |  |
| Favored (%) | 98.21% | 95.64% |
| Allowed (%) | 1.68% | 4.25% |
| Disallowed (%) | 0 | 0 |

**Extended Data Table 2.** Potencies and efficacies of DAMGO in MOR signaling assays using wildtype and mutant MOR in the presence and absence of different BMS-986187 concentrations. Data represent mean EC50 or pEC50 ± SEM, efficacy Emax as Span ± SEM, and ΔEmin ± SEM (calculated as Emin_BMS-986187_-Emin_DMSO_). Biological repeats for each dataset are indicated, and all experiments were performed in triplicate.

**cAMP Accumulation**
| MOR Construct | Replicates |  | DMSO |  |  | DAMGO |  |  | BMS-986187 |  |
| --- | --- | --- | --- | --- | --- | --- | --- | --- | --- | --- |
|  |  |  | EC50 (nM)<br>pEC50±SEM | Span ± SEM (%) |  | EC50 (nM)<br>pEC50±SEM | Span ± SEM (%) |  | EC50 (nM)<br>pEC50±SEM | Span ± SEM (%) |
| WT | 3 |  | ND | ND |  | 3.50<br>8.36±0.20 | 100.00±0.00 |  | ND | ND |

**cAMP Accumulation**
| MOR Construct | Replicates |  | DMSO |  | 3 uM BMS-986187 |  |  | 10 uM BMS-986187 |  |
| --- | --- | --- | --- | --- | --- | --- | --- | --- | --- |
|  |  |  | EC50 (nM)<br>pEC50±SEM |  | EC50 (nM)<br>pEC50±SEM | ΔEmin<br>(span ± SEM %) |  | EC50 (nM)<br>pEC50 ± SEM | ΔEmin<br>(span ± SEM %) |
| WT | 3 |  | 5.00<br>8.27±0.22 |  | 1.26<br>8.83±0.06 | 12.55±4.37 |  | 0.55<br>9.22±0.06 | 27.14±6.52 |
| W194A | 4 |  | 1.74<br>8.74±0.11 |  | NA | 70.26±4.41 |  | NA | 74.42±2.11 |
| M205A | 4 |  | 7.45<br>8.22±0.21 |  | 6.40<br>8.24±0.13 | -1.86±5.07 |  | 3.99<br>8.40±0.10 | 1.5±8.4 |
| Y151L | 4 |  | 0.36<br>9.55±0.22 |  | 0.53<br>9.36±0.16 | 1.54±7.91 |  | 0.45<br>9.34±0.05 | 2.58±5.21 |
| Y151A | 3 |  | 0.17<br>9.85±0.16 |  | 0.24<br>9.77±0.36 | 9.65±7.65 |  | 0.26<br>9.68±0.22 | 5.47±4.83 |
| Y151F | 3 |  | 1.33<br>8.94±0.08 |  | 0.21<br>9.81±0.23 | 29.76±4.48 |  | 0.18<br>10.02±0.26 | 53.96±4.52 |
| L118A | 4 |  | 3.20<br>8.58±0.09 |  | 1.14<br>9.11±0.23 | 9.76±2.98 |  | 0.57<br>9.42±0.09 | 27.61±10.13 |
| I144A | 5 |  | 1.38<br>8.90±0.09 |  | 0.29<br>9.61±0.26 | 5.36±1.41 |  | 0.18<br>9.9±0.26 | 32.95±4.8 |
| I148A | 5 |  | 0.15<br>9.94±0.13 |  | 0.04<br>10.48±0.25 | 10.55±3.7 |  | 0.05<br>10.52±0.30 | 39.61±4.24 |
| S198A | 3 |  | 0.20<br>9.75±0.18 |  | 0.27<br>9.64±0.17 | 14.02±5.08 |  | 0.20<br>9.89±0.24 | 29.59±1.79 |

**Gi1 BRET**
| MOR Construct | Replicates |  | DMSO |  | 3 uM BMS-986187 |  |  | 10 uM BMS-986187 |  |
| --- | --- | --- | --- | --- | --- | --- | --- | --- | --- |
|  |  |  | EC50 (nM)<br>pEC50±SEM |  | EC50 (nM)<br>pEC50±SEM | Span ± SEM (%) |  | EC50 (nM)<br>pEC50±SEM | Span ± SEM (%) |
| WT | 3 |  | 16.26<br>7.84±0.25 |  | 4.20<br>8.24±0.28 | 96.74±2.96 |  | 1.39<br>8.85±0.15 | 94.30±4.63 |

**Extended Data Table 3.** Composition of the MD simulation boxes.

|  | BMS986187-DAMGO-bound<br>MOR-G<sub>I1</sub> | DAMGO-bound MOR-G<sub>I1</sub> |
| --- | --- | --- |
| Simulation Box Dimensions (Å) | 160x157x152 | 161x154x147 |
| Total Number of Atoms | 267,595 | 258,151 |
| Water Molecules | 54,865 | 52,402 |
| Salt Concentration | 0.15M | 0.15M |
| Cholesterol | 166 | 171 |
| POPC | 181 | 167 |
| POPE | 189 | 178 |
| POPS | 54 | 54 |
| PIP2 | 38 | 37 |
| PSM | 51 | 58 |
| GM3 | 35 | 36 |

